# Optimal Time Lags from Causal Prediction Model Help Stratify and Forecast Nervous System Pathology

**DOI:** 10.1101/2021.03.05.434171

**Authors:** Theodoros Bermperidis, Richa Rai, Jihye Ryu, Damiano Zanotto, Sunil K Agrawal, Anil K. Lalwani, Elizabeth B Torres

## Abstract

Traditional clinical approaches diagnose disorders of the nervous system using standardized observational criteria. Although aiming for homogeneity of symptoms, this method often results in highly heterogeneous disorders. A standing question thus is how to automatically stratify a given random cohort of the population, such that treatment can be better tailored to each cluster’s symptoms, and severity of any given group forecasted to provide neuroprotective therapies. In this work we introduce new methods to automatically stratify a random cohort of the population composed of healthy controls of different ages and patients with different disorders of the nervous systems. Using a simple walking task and measuring micro-fluctuations in their biorhythmic motions, we combine non-linear causal network connectivity analyses in the temporal and frequency domains with stochastic mapping. The methods define a new type of internal motor timings. These are amenable to create personalized clinical interventions tailored to self-emerging clusters signaling fundamentally different types of gait pathologies. We frame our results using the principle of reafference and operationalize them using causal prediction, thus renovating the theory of internal models for the study of neuromotor control.

## Introduction

Patients with Fragile-X Tremor Ataxia syndrome (FXTAS), patients with Parkinson’s Disease (PPD) and patients in the broad spectrum of Autism (ASD) my all have motor issues detectable and tractable through gait patterns at different stages of their lifespan ^1–8^. Generally, these disorders share some common genes ^9–11^, but their phenotypic features differ depending on the person’s age and on the evolution of various epigenetic factors ^1,12.^While the autistic patient may arrive at the diagnosis via Psychiatric/Psychological measures, devoid of motor criteria, it is highly probable that the PPD will do so through motor criteria from a movement disorders specialist in Neurology. In contrast, the FXTAS patient may receive an ASD diagnosis during childhood, when differences in communication and social interactions gain priority over motor issues ^13,14^ but then, later in life, as motor issues exacerbate, they may gain the diagnosis of PPD. FMR1-premutation carriers may be considered neurotypical until later in adult life, when tremor, ataxia, and symptoms of parkinsonism become highly visible ^15,16.^Indeed, the FMR1 CGG repeat length predicts motor dysfunction ^17^ of the type that PPD manifest much later in life.

We know that motor activity reveals that aging with Autism greatly departs from typical aging ^18^ despite whether the person is or is not on psychotropic medications ^19^. We are also aware that parkinsonism, typically associated with human aging ^3,20,^is prevalent in Autistic adults ^21^. Indeed, the rate of parkinsonism in the general population aged 65–70 has been estimated at 0.9 % ^20^, compared to 20 % in ASD aged after 39 ^21^, when excess involuntary motions at rest have been quantified ^18,22.^

Parkinsonism in FMR1-premutation carriers may be indistinguishable from PD ^23^. And Fragile-X syndrome (FXS) has high penetrance in Autism ^24^ with genetic overlap across Parkinson, Ataxias, ASD and FXTAS ^9^. Furthermore, in Autisms of known etiology, such as SHANK3 deletion syndrome ^25^, a constellation of sensory and cognitive issues parallels motor abnormalities detectable through their gait ^8^. The genetic and clinical overlap across these disorders motivate us to search for similarities and differences in relation to a well-known disorder of gait. Using the screening version of the vestibular Dizziness Handicap Inventory (DHI-S) ^26,27^ we compare the neuromotor dynamics of gait in patients ranked according to this clinical scale ^28^, to patients with the above-mentioned clinical disorders. We posit that if we were to automatically stratify a random draw of the population containing these disorders (alongside age- and sex-matched controls) we may be able to eventually tailor gait-related accommodations to highly heterogeneous disorders such as ASD. Since there is a common pool of genes affecting both neurodevelopment and neurodegeneration, identifying clinically-characterized gait issues in these disorders, might open a new path to pre-emptively offer neuroprotective motor-sensory-based accommodations in early neurodevelopment gone astray ^9^.

Gait studies integrated with wearable sensors in natural activities have been very useful to ascertain various aspects of these phenotypes ^16,29–31,^ but population statistics employed in these analyses are not personalized. They primarily focus on descriptive summary statistics from a priori assumed distributions, a procedure that incurs in loss of gross data ^32^. Further, owing to anatomical disparities confounding gait parameters, we need scaling methods that standardize outcomes and provide proper similarity metrics to compare across different subtypes of a given disorder. There is room to examine individual motor fluctuations within the context of dynamic network analyses and causal prediction ^33^. Here we integrate causal and non-causal network connectivity analytics to examine self-emergent patterns from standardized micro-movement spikes derived from gait-kinematics. We discuss our results within a new unifying framework to study disorders of the nervous systems across different diagnoses of disparate clinical criteria and yet, converging functional phenotypes.

## Results

Granger causality (GC) is a mathematical concept that allows us to quantify the causal effect of a stochastic process A on a second stochastic process B. It compares the prediction error of a dynamic model that predicts the evolution of B without the presence of A with the error of a prediction model that includes both A and B. If the inclusion of A improves the prediction, we say that A causally predicts B (A**→**B) and we can assess the degree to which this is determined by the method of choice. Our approach to modeling and calculating the Granger causality assumes models with a single internal delay. This allows us to find, within a particular range, the internal delay that maximizes the causality from A to B. To implement our methods using time series data from a grid of 23 sensors collecting bodily orientations and acceleration as the person walks, we first extract the angular speed time series (rad/s) and the linear accelerations (m/s^2^) over the span of 3-min walks. We then, build a data type that we have created, the micro-movements spikes (MMS ^34^) which extracts the moment-to-moment fluctuations in the signal’s amplitude provided by the peaks, and the fluctuations in inter-peak interval timings. This data derived from the sensors is unitless as it is standardized to characterize the dynamic behavior of motion by scaling out anatomical differences. This standardization of the data also enables us to compare the stochastic signatures across these participants’ motions in a random draw of the population, while considering different ages and different clinical genotypes and phenotypes. As we keep the indexes into the original physical unit ranges, we can further estimate cross-population scales and boundaries for each parameter of interest.

Upon standardization, we apply our technique between each directed pair of MMS time series across all 23 nodes corresponding to different joints across the full body. The maximum outwards GC of each node towards the rest of the body for the average GC-directed network of each group revealed marked differences from group to group. Furthermore, the average Optimal Lag network of maximum lags from a node to the rest of the body, assuming that the MMS series of that node, is the causal stochastic process, also revealed fundamental differences on timing across the different ages and nervous systems’ disorders. We detail them below:

### Granger-Causal Networks Reveal Age-Dependent Gait Differences in Typical Controls

As age increases in healthy participants, we see fundamental differences emerging across the patterns of GC prediction between their upper and lower body. The relative causal connectivity between upper to lower body is higher in elderly participants than in young controls. This can be appreciated in Figure 1A using an anthropomorphic interconnected network representation of the full body using the 23 nodes. The size of the node is proportional to the GC out degree value of that node as it connects to all other nodes. Furthermore, elderly control participants have in general lower optimal lag times of the lower extremities (*p*=0.0293, *t-test*) suggesting an overall slowdown in information transmission and feedback between upper and lower extremities. Often, patterns such as bradykinesia and loss of gait control are found in pathologies of the nervous system. Yet, here we observe them as a part of the natural aging process in this cross-sectional cohort of healthy controls. We use a colormap representation to indicate the lag values in Figure 1A, whereas in Figure 1B, we provide the empirical frequency histograms of the interpeak interval timings of the MMS. The corresponding physical times expressed in seconds distribute according to the continuous Gamma family of probability distributions, optimally fit to the empirical data, in a maximum likelihood estimation (MLE) sense. These patterns are depicted for the healthy controls in Figure 1B (top panels) while empirically estimated summary statistics are depicted in Figure 1C.

**Figure 1:**
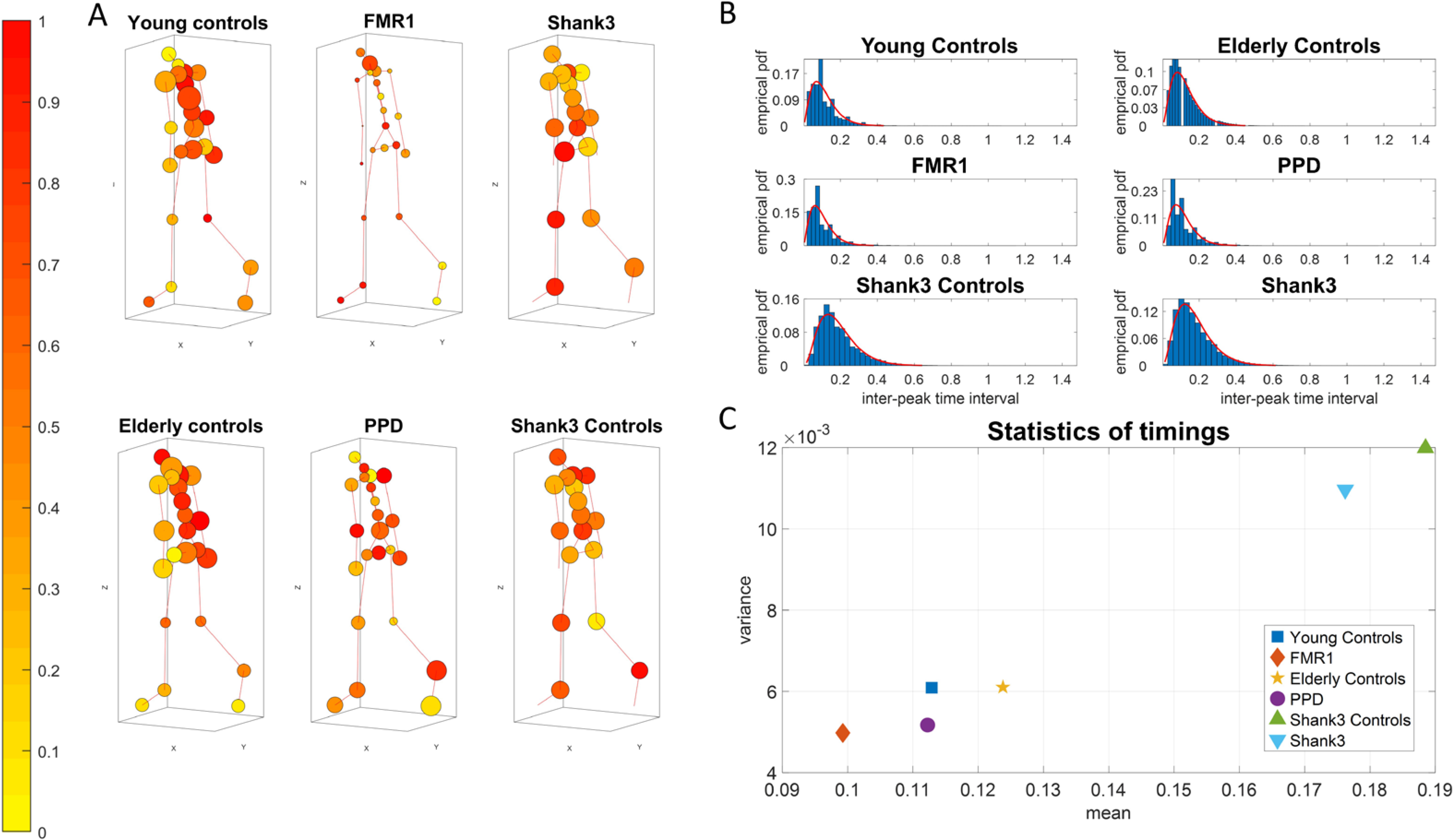
GC Optimal Lag Maps and timing information. **A)** Anthropomorphic network representation of human walking patterns. The nodes’ size is proportional to the number of outgoing links connected to the other nodes in the network (out degree) of the average GC network. The color of the nodes is the maximum optimal lag response of a node with the rest of the body for the average Lag Response network. A unit of one lag refers to the distance in time between two peaks. The actual physical time between the two peaks can vary. **B)** The frequency histograms depict the distribution of the inter-peak timings for each group. The empirical distributions are well fitted by the gamma family of distributions. **C)** Large departure from ideal patterns at college age are observed in the youngest developing (still growing) children and in the elderly controls. Further separation of the various pathologies is captured in relation to age-matched controls. For the youngest controls (5-7 years old) from the SHANK3 dataset, the average inter-peak timing is 0.1885 seconds with a std of 0.1095, compared to the SHANK3 participants 0.1762 with a std of 0.1047. For the young controls of college age, 0.1129 with a std of 0.0780. For the elderly controls, 0.1238 with a std of 0.0781. For the FMR1 carriers, 0.0993 with a std of 0.0705. For the PPDs, 0.1122 with a std of 0.0719.

### FMR1-Carriers are Different from Controls of Comparable Age and from PPD

Premutation carriers of the FMR1 gene show a significant decrease in GC connectivity across the body in relation to the young controls of comparable age (*p*=0.0061; *t-test*) This is depicted in Figure 1A along with fundamental differences in the patterns of optimal lags, which are abnormally lower than controls (*p*=2.7160e-04; *t-test*). This reduction in optimal lag values in the premutation carriers is accompanied by a reduction in their timing variability that resembles (and perhaps forecasts) the type of reduction in variability observed in the PPD. The latter shows a significant reduction in GC across the body relative to the elderly controls (*p=0.0021; t-test*). This is particularly the case in the torso, with lower lags in the lower body, when we compare them to the elderly controls. Figure 1C summarizes the statistics of the timing. This parameter space reveals healthy patterns of timing variability in the controls *vs.* atypically lower variability patterns in the premutation carriers and the PPD.

### Motor Timings and Inter-peak Interval Timings Differ Between SHANK3 Participants and Age- and Sex-Matched Controls

Subjects diagnosed with SHANK3 deletion syndrome exhibit overall higher motor timings, especially in the upper body nodes, compared to age- and sex-matched controls, but also in relation to patients with other disorders. This can be appreciated in Figure 1A-C, where we express the differences across an avatar representation (1A), using all nodes accessible to our instrumentation, frequency histograms of interpeak interval timings fitted by empirical *probability density functions (*PDFs) (1B) and a parameter space (1C) summarizing the statistics derived from the average empir*i*cally estimated behavior of the cohort. Regarding inter-peak interval timings from the derived Micro-Movement time series of all nodes, SHANK3 *deletion participants* show significantly lower mean values from the*ir age- and sex-matched* controls, following a similar pattern to that observed in FMR1 carriers. SHANK3 deletion participants are often diagnosed *with* ASD, as are some FMR1 premutation carriers. The two groups here exhibit lower inter-peak interval timings than age- and sex-matched neurotypicals controls who performed the gait task under the same settings.

### Prevelance of Causal Feedback Loops is Much Higher in the Controls

An important concept in the theory of neuromotor control, first introduced by von Holst and Mittelstaedt in 1950, is known as the principle of reafference ^35^. Every time a motor movement is to be initiated, a motor signal is sent to the periphery to perform a certain movement through the efferent nerves, which is called efference. Similarly, the sensory input coming from the periphery towards the Central Nervous System (CNS) is called afference. The afference consists of two components. The first component is called exo-afference and is the afferent input generated from the environment. The second component is called endo-afference, which is the sensory input self-generated from the body’s own actions. We discovered that during complex actions, the endo-afference is separable into intended segments (under voluntary control), and incidental segments (occurring spontaneously and largely beneath awareness ^36^.) Because of its internal origins and its relation to movements, the latter is known as kinesthetic reafference, though it also includes pain and temperature afference from the corresponding fibers at the periphery. According to the principle of reafference, every time a movement is initiated by sending information to the motor system, a copy of the signal is created, called the efferent copy or corollary discharge, and is sent to the CNS to inform of the impending movement. This enables the CNS to distinguish sensory signals stemming from external environmental factors, from sensory signals coming from the body’s own actions. The efference copy is provided as input to a forward internal model to predict the sensory consequences of the motor action to be initiated. Comparing the predicted movement with the actual movement is precisely what allows the CNS to recognize its own actions and at a higher cognitive level, form a sense of self. In the internal models of neuromotor control, the intended consequences from voluntary acts can be evaluated using the principle of reafference ^37^. A non-trivial extension of this idea is to include, in addition to consequences from voluntary actions, the unintended consequences from those incidental actions that occur spontaneously, without instructions or precise targets. This idea has been proposed and investigated by our group, to define different levels of volition ^36,38–40,^ and offered as a fundamental ingredient of the more general concept of motor agency.

The principle of reafference has been a source of inspiration for decades in biology, cognitive science, vision, neuromotor control and robotics, and attempts to model it fall usually within the realm of control theory and dynamical systems ^41^. Our approach to operationalize this principle does not assume a particular deterministic feedforward model. It rather assumes stochastic dynamic models for each body node and for pairs of body nodes. If the principle of reafference implies the presence of feedback loops between the CNS and the periphery, the fact that information is carried along efferent and afferent pathways and processed in various organs of the CNS implies that it is not an instantaneous process, but it is characterized by some internal timing. In fact, timing is important to accurately predict the consequences of one’s actions, coordinating such actions in space and time. Timing plays a central role in agency, leading to a sense of action ownership and self-awareness. Hence, the causal analytics, being sensitive to the choice of timing of the model, can be used to estimate the actual motor timings in the nervous system.

But how exactly can we detect the presence or not of a feedback loop between two nodes? Once again, the answer can be found in one of Granger’s definitions in his original paper ^42^. Following the formal definition of causality, he proceeds to define feedback. According to the definition, if A and B are two stochastic processes, and A causes B in the Granger sense, but also B causes A in the Granger sense, *i.e.*, causality occurs in both directions, then that implies the presence of a feedback loop between the two processes. Then, simply checking for each undirected pair of body nodes, we can check whether the causal connectivity graph that we derived earlier exhibits causality in both directions. Then, we can determine whether there is a feedback loop between the nodes, within the accuracy of our model and the choice of timing.

Addressing Granger’s notion of feedback (up to our limited assumption of an autore*g*ressive model explained in the Materials and Methods Figure 14) provides the results depicted in Figure 2. As the results from Figure 1 suggested, the healthy controls show (cross sectionally) the effect of natural aging, this time, on the feedback patterns that we take pairwise across these 23 nodes of the body. Consistent with the Figure 1A summarizing the group patterns (in one direction A**→**B), we see the patterns of feedback loops across each group, detected when including B**→**A and obtaining GC in both directions. Young healthy controls have a more distributed pattern across the body, in contrast to elderly controls, who show a diminished pattern in the lower extremities accompanied by an abnormally higher feedback-loop pattern across the upper extremities, perhaps compensating for their lacking in the lower extremities. This decrease in feedback loops generalizes across all bodily nodes in PPD, but interestingly, the decrease is even more evident in the FMR1 premutation carriers, closer in age to the healthy young controls. At such a young age compared to the PPD, these patterns in the FMR1 premutation carriers forecast trouble on the horizon, of the type that elderly PPD eventually experience. The younger controls (for the SHANK3) show fewer feedback loops than the older controls (for the FMR1/PPD group), a result that might be explained as the older participants have reached full gait maturation, but the young ones are still growing and adapting their gait. Participants with the SHANK3 deletion have a higher prevalence of feedback loops than those age-matched neurotypicals. This is consistent with lack of synergies across their body ^8^, a feature that makes the brain’s motor control problem rather challenging, as with the FMR1 pre-mutation carriers. Both disorders have high ASD penetrance, as does the prevalence of their gait issues across the lifespan. Further differentiating across these groups will be important to refine gait pathologies and their possible origins in neurodevelopmental *vs*. neurodegenerative stages. This outcome pertaining their neuromotor dynamics reveals important neurological characteristics of each disorder.

**Figure 2:**
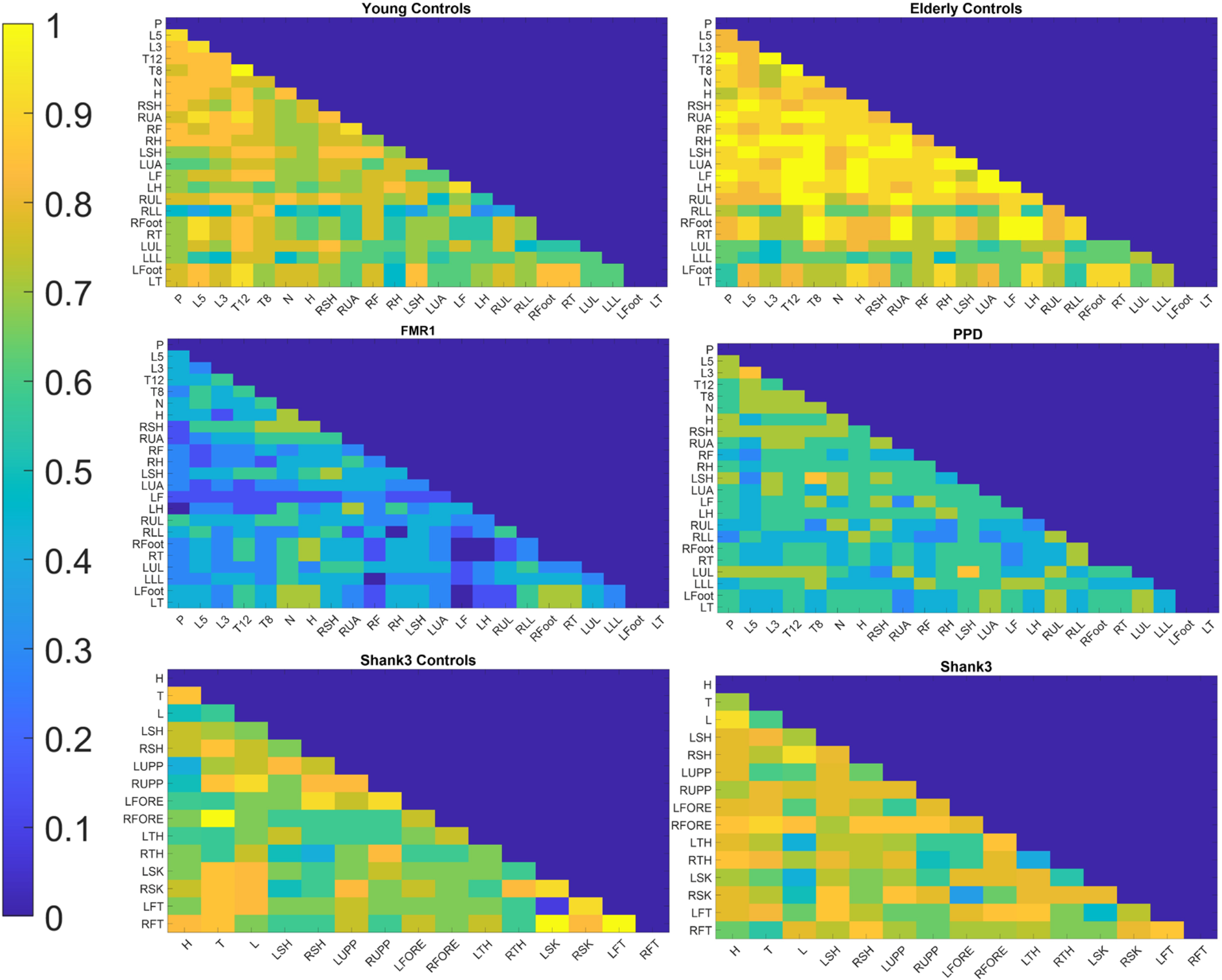
Feedback loops differ across groups. For each group, the figures show the percentage of subjects within the group for whom a feedback loop was detected between each pair of body nodes. Young controls (of college age) differ from the elderly controls who, on average, show higher feedback loops in the upper body, whereas some very young controls (here called SHANK3 controls) 5-7 years of age are still maturing their gait, and developing modular synergies across the body. They differ from the SHANK3 deletion syndrome, who show lack of synergies simplifying the motor control problem that abundant degrees of freedom pose to the brain. Patients with Parkinson’s disease show an atypical lower feedback across the body, which is further exacerbated in FRM1 premutation carriers, despite their youth.

### Identification of Different Groups’ Subtypes

One of the current challenges in data-driven analyses is to automatically identify self-emerging subtypes of a disorder on a spectrum, as patterns stratify to form clusters of data points across a given scatter. We here address this automatic stratification of a random draw of the population composed of different types of patients and ages, using various parameter spaces derived from the stochastic patterns of the gross data that is often thrown away or smoothed out through grand averaging.

For each participant from the FMR1/PPD dataset, we calculated the trajectory of the Center of Mass of the lower body in 3D space and we extracted the MMS series from the Euclidean norm of the position time series. Upon obtaining the standardized (scaled) speed amplitudes, we obtained the peaks and calculated the empirical distributions of the inter-peak-interval timings of these MMS series. We noted the presence of multiple modes in the frequency histograms and performed the Hartigan Dip test of unimodality ^43^. We performed this statistical test both on the patterns from each participant separately and pooling the data across participants for each group. For the latter, we did so after concatenating the MMS series data of all subjects within a group. A similar analysis was performed for each participant from the SHANK3/DHI dataset. The Hartigan Dip Unimodality test per group showed that for all groups the total empirical distributions fail the unimodality test. All groups distributed bimodally, yet how the modes grouped had specific patterns particular to each group. This is depicted in Figure 3.

**Figure 3:**
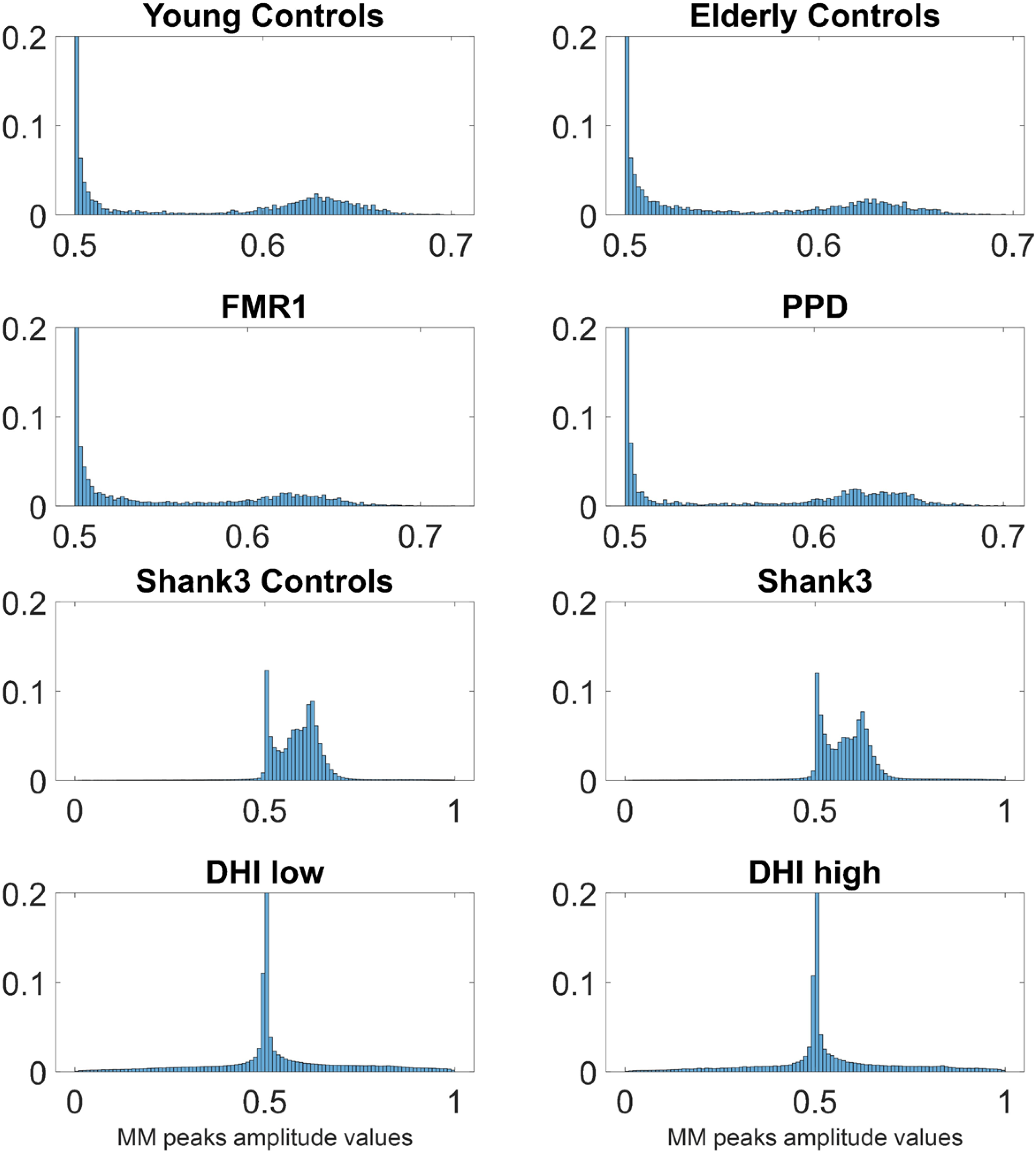
Unimodality *vs.* Multi-modality in the distributions of MMS from peak angular speed amplitudes differentiate the various groups. Different modes are specific to each cohort, mode 1 exponential and mode 2 Gaussian, with different dispersions and percentages of points in each mode. In contrast to other groups, both the low- and the high-score DHI groups, exhibit unimodal (highly kurtotic) distributions.

The first mode with lower values of the standardized MMS peaks amplitude was well fit by an exponential distribution, whereas the second mode was well fit by a Gaussian distribution that had the lowest dispersion in healthy young controls. In elderly controls, the percentage of points in the exponential mode increased, as did in FMR1 premutation carriers (closer in age to the young controls) and the PPD, with the latter showing distribution patterns comparable to the carriers. To see this, for each participant, we found in which mode the MMS activity was highest and calculated the percentages of participants per group with the highest activity in each mode. Most young and elderly controls have the highest MMS activity in the Gaussian mode. FMR1 carriers and PPDs have higher prevalence of activity in the exponential mode. Furthermore, the SHANK3 deletion participants have higher prevalence of activity in the exponential mode, compared to their age- and sex-matched controls. Moreover, SHANK3 controls have the lowest exponential mode activity out of all groups, a result that may reflect the early maturational stage of their developing nervous system in a rapidly growing body (Figure 4.) This result suggests that healthy gait requires a proper balance between these two modes, and that examination of their distribution across the population could provide clues on the status of the person’s gait health.

**Figure 4:**
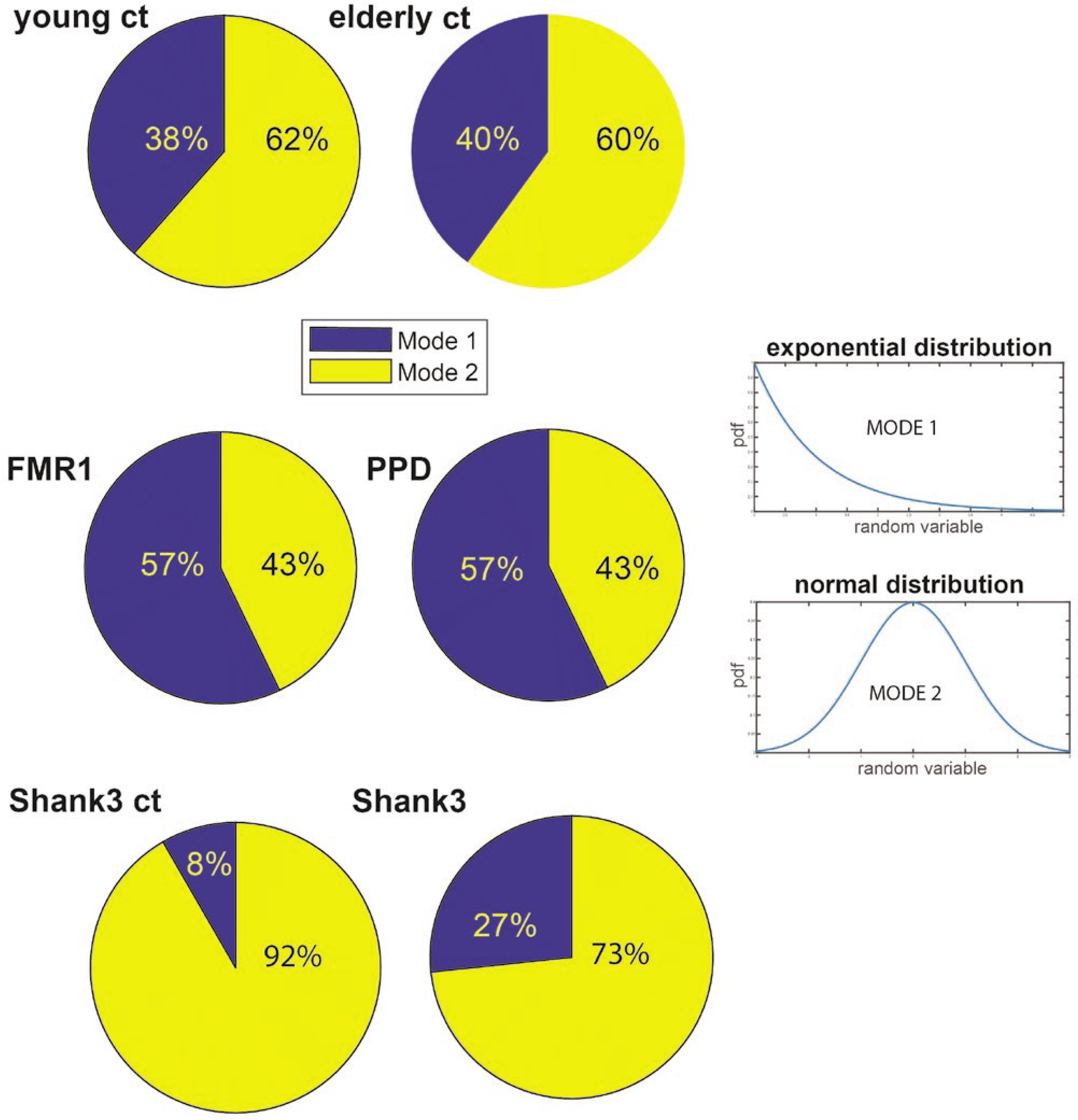
Percentages of distribution’s modes varies across groups with lowest Exponential mode in young controls and highest in FMR1 carriers and PPD. Gaussian mode is highest in the young and elderly controls and comparable between PPD (higher) and FMR1-carriers. Gaussian mode is significantly higher in participants with SHANK3 deletion syndrome, large departure from the youngest controls in the study. The exponential mode is at the lowest in very young developing children, higher in college age controls and at its highest in the elderly controls. An excess in this mode suggests pathological gait patterns in FMR1 and PPD.

These patterns motivated a parameter space whereby we represent points as triplets-points in a three-dimensional space, with coordinates representing the % of Mode 1 (exponential), the Dip value from the Hartigan’s Dip Unimodality test, and a scalar, giving the speed to frequency ratio (we discuss below the motivation for using this ratio.) We then performed k-means clustering analyses on the scatter of points from each group, to identify potential subtypes of activity in each cohort of participants. Figure 5A depicts the results (under the Euclidean distance metric; note that using the first mode in this representation is sufficient, as the second mode percentages are complementary, *i.e.,* the sum of percentages add to unit). Here the young controls have 3 clusters, and the other groups show 2 clusters (cluster elements are represented proportional to the size of the marker.)

**Figure 5:**
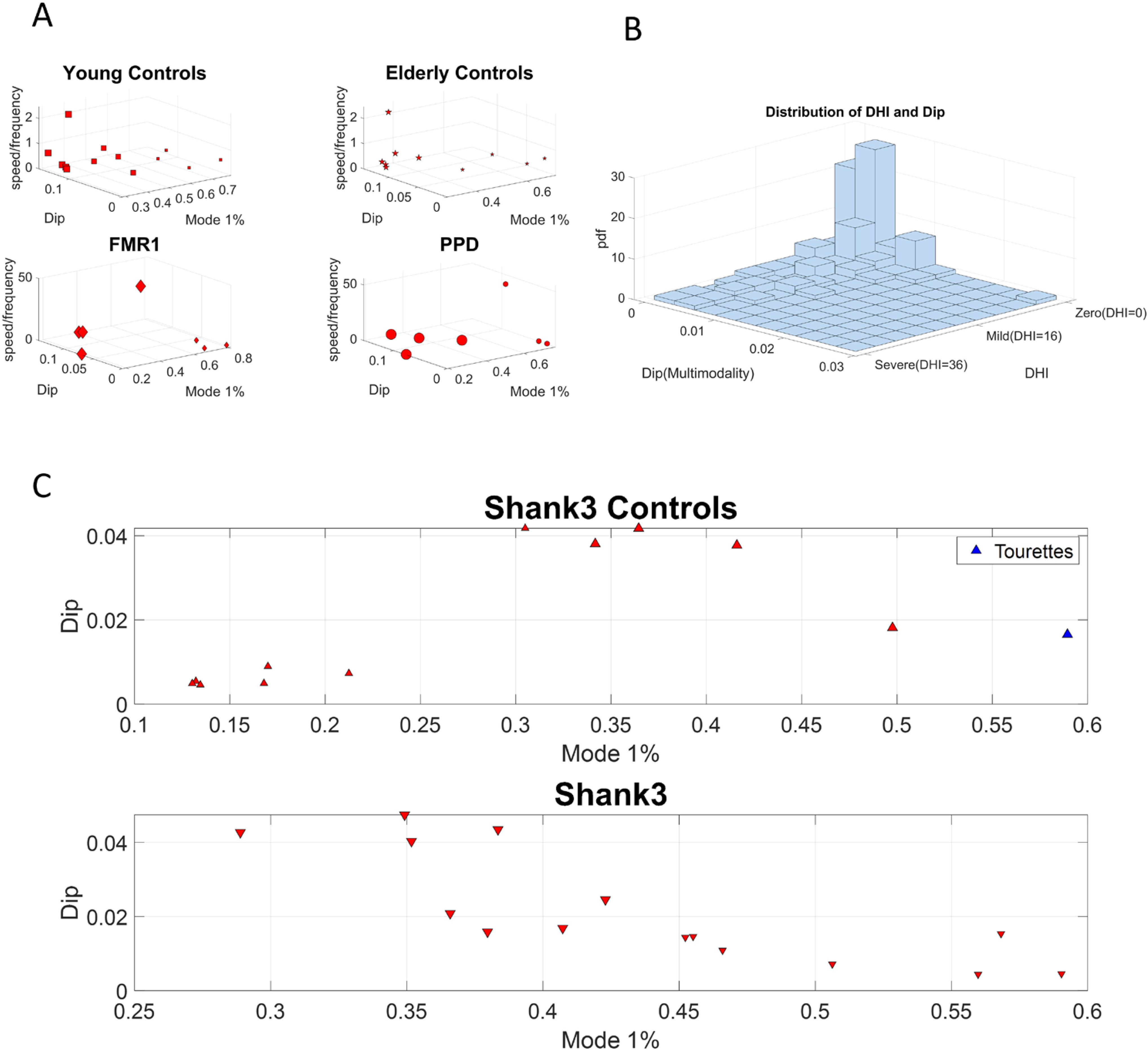
Subgroups within each group revealed by group’s clusters. **A, B)** The clusters for each group using the k-means clustering algorithm with 3 for young controls and 2 for each of the other groups. Subtypes of activity emerge for all groups. The empirical distributions for the MMS of the norm of the lower body center of mass, COM 3D position or lower body angular acceleration (for subjects from the SHANK3 study) data series for the 6 groups are bimodal for all 6 groups. **B)** For subjects in the DHI study, we calculated the empirical bivariate distribution of DHI scores and dips. **C)** For the controls from the SHANK3 study low exponential mode activity implies high dip values. For the SHANK3 participants, high exponential mode activity implies low dip values. Note that a participant with Tourette’s syndrome (manifesting an excess of motor ticks) and who also received an ASD diagnosis, had the maximum exponential activity over all subjects.

We identified 3 sub-types for the young controls group and 2 sub-types for the elderly controls. There is an apparent negative correlation between higher dips and lower first mode (exponential) activity. The FMR1 carrier group has two subtypes. The first subtype has high first mode activity and low dip and the second subtype has low first mode activity and high dip. A similar subtyping pattern is observed for the PD group. SHANK3 controls exhibit a positive correlation between exponential mode activity and dip, whereas participants with SHANK3 deletion syndrome exhibit a negative correlation between those two. We localized an ASD participant with Tourette’s syndrome in this space, to examine high levels of exponential mode activity (the maximum of the cohort) considering the excess motor ticks of this participant. This localization helps us interpret the result as it sets bounds on the parameter space for cases of ASD of known etiology. Furthermore, the bivariate empirical distribution of DHI scores and dip values showed that the vast majority of DHI participants follow unimodal distributions of motor activity, while any multimodal activity is detected mostly in mild to moderate DHI score individuals. The results from the distribution analysis can be appreciated in Figure 5B. In 5A and 5C the size of the marker denotes the group within each cohort. Differences in unimodality across DHI participants with severe scores, combined with differential patterns in MMS peak amplitude values in Figure 3 hint at possible different groupings, a question that we further explore below.

### Groups Differ Relative to Young Controls within a Parameter Space from the Data Stochastic Features

The above results motivated us to quantify the departure of each group from the young controls of college age (as an ideal, normative state of motor control.) To that end, since the lower body and the pelvis play a crucial role in gait control, we first characterized the patterns of fluctuation in the motions of the pelvic area and upper leg nodes) by fitting the continuous Gamma family of probability distributions as the optimal family in an MLE sense. Through this process, we obtained the scale parameter, which in the case of the Gamma family, is also the noise to signal ratio (NSR.) We obtained the average NSR and computed skewness of the MMS series of the pelvic area for each subject. We also obtained the GC network asymmetry for each subject (See Methods). This paramer space is shown in Figure 6 across all three comparisons.Two sample *t-tests*, showed that young controls differ from elderly controls in mean skewness (with bortherline significance, *p-value=0.096*, *confidence interval=[0.0465, 0.0299]*). They also differ from PPDs, signficantly at the *0.05 alpha level*, in mean pelvic NSR (*p-value=0.0193*, *confidence interval=[−0.047, −0.00047187])*. Finally, they differ significantly from FMR1 carriers in assymetry *(p-value=0.0107, confidence interval=[−0.3386, −0.051]).* In each plot, we fitted surfaces over the data samples for the young controls to help visualize the differences and localize each cluster shifting across the parameter space relative to this normative surface.

**Figure 6:**
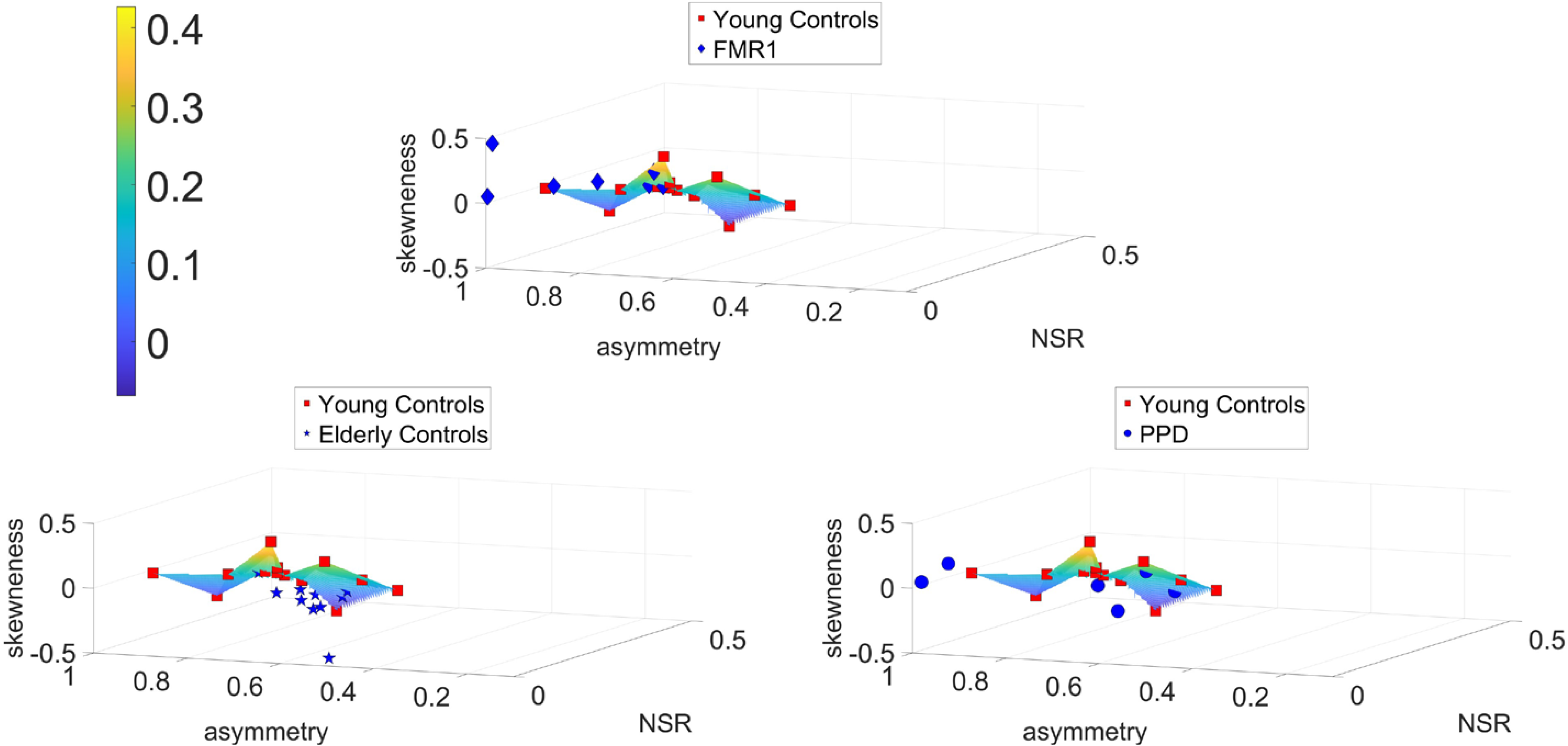
Parameter space reveal differences across controls, elderly controls and PPD participants. NSR, GC network asymmetry and average Skewness are used to build a parameter space showing departure from ideal young controls’ scatter fitted by a surface. Scatters from the FMR1carriers, the elderly controls and the PPD largely depart from young controls’ surface. Color bar reflects skewness range of values.

Notice in Figure 6 that FMR1 and PPD depart similarly from the normative surface of young controls, while the elderly controls occupy a different location in this parameter space.

Normalized causality, optimal timings, and efferent-afferent asymmetries in the causal and motor timing activity of subjects from the SHANK3 deletion syndrome dataset separated them from the corresponding age- and sex-matched SHANK3 controls. This separation had a pattern analogous to the case of FMR1 participants, showing a consistent departure in causal dynamics from the corresponding neurotypical cases. The subject diagnosed with Tourette’s syndrome showed extreme values across all parameter spaces. His motor patterns included excessive motor ticks that visibly interfered with his gait. This extreme case helped us interpret those high bounds in exponential mode depicted in Figure 5. The results of the new analysis can be appreciated in Figure 7.

**Figure 7:**
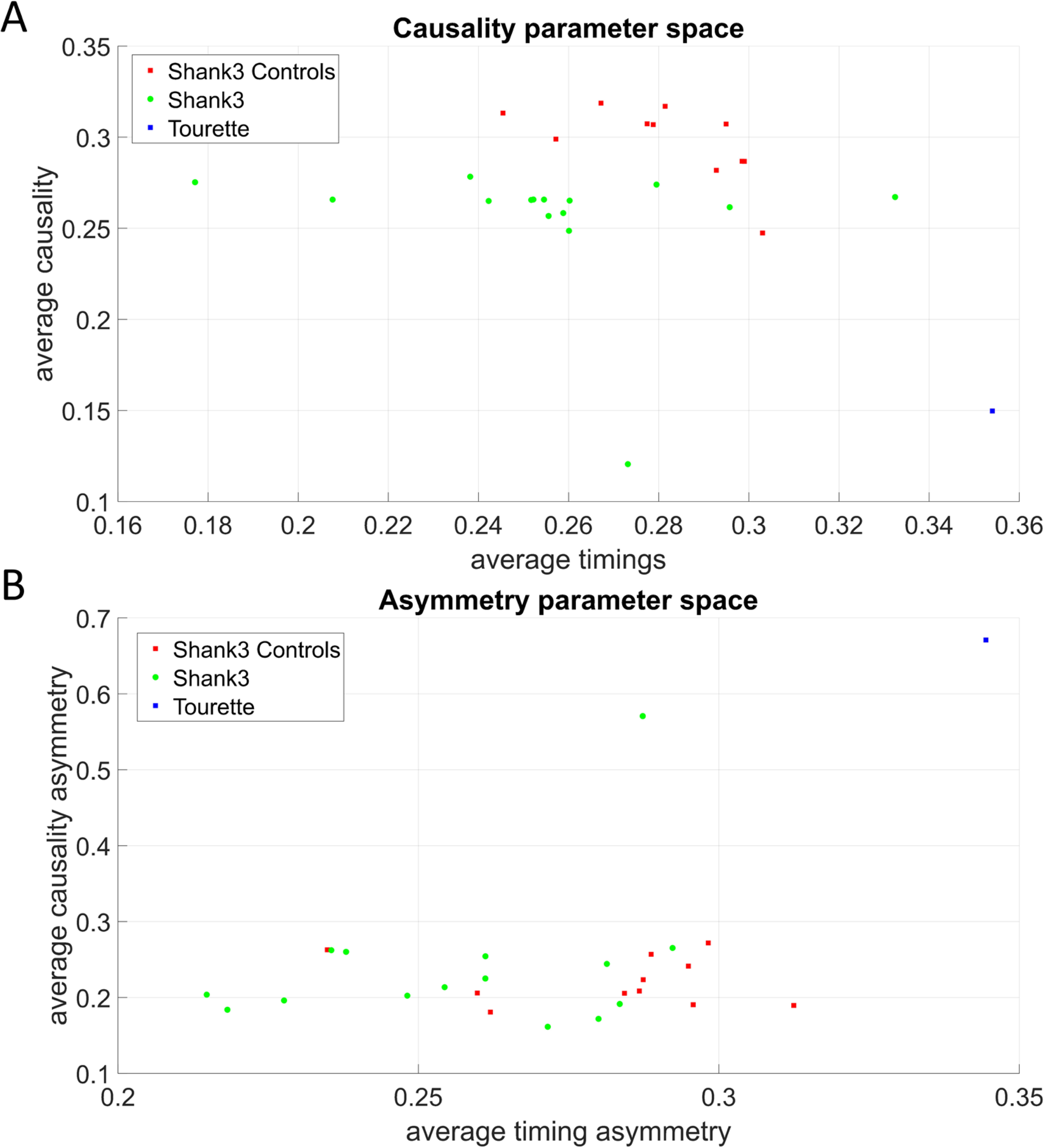
Parameter spaces reveal differences across ASD-SHANK3 deletion participants and the youngest controls of the study. **A)** Normalized values of average causality and motor timings separates these participants with ASD of known etiology from age- and sex-matched controls. The one control subject diagnosed with ASD Tourette’s syndrome is an outlier and has the highest average motor timings value. **B)** Normalized values of timings asymmetry also separate this type of ASD from controls and once again, the ASD-Tourette’s syndrome participant is an outlier that exhibits the maximum value of asymmetry both in causality and timings.

### Automatic Stratification of a Random Draw of the Population Suffering from Gait Disturbances Distinguishes Between Gait Pathologies from Vestibular-Cortical and Sub-Cortical Problems

We considered 118 participants with gait disturbances involving dizziness, the sensation of vertigo, fatigue, presyncope and disequilibrium. These are quantified through the vestibular handicap inventory, the Dizziness Handicap Inventory screening tool (DHI-S) ^28^. This tool spans scores ranging from zero-value, with a cutoff at 16, whereby zero-16 are considered mild, while those above 16 are considered severe. We used these scores in conjunction with the digital data from both feet as participants walked in similar ways as those suffering from other disorders and typically developing or aging. Using these DHI ranges, we then examined the cohort of controls across the lifespan, as well as the cohorts of PPD, FMR1 and those with a diagnosis of ASD of known etiology (SHANK3 deletion syndrome) and of idiopathic ASD. Our motivation here was to combine the MMS data type, our methods of analyses and various parameter spaces across all 187 participants, to localize various neurological disorders and ASD, with an eye for self-emerging clusters. We seek to characterize these disorders in relation to gait from vestibular handicap, to ask if the gait issues that we detected in ASD participants are closer to those quantified in DHI elderly, or to those found in the PPD elderly or the FMR1 participants. Unlike the vestibular handicap gait issues, the latter are linked to subcortical neurodegeneration rather than higher level sensory perception and integration. Given the current ASD definition relying on higher-level perceptual and sensory issues, and the high prevalence of ear infection and auditory brain stem issues since early infancy ^44^, it is possible that their gait problems align with the vestibular integration, rather than the subcortical motor control problems. This hypothesis is addressed below:

First, we represent the DHI participants in a normalized parameter space spanned by feet causal forces (defined by the ratio of causality divided by the corresponding time lags), and average timings. We then identify fundamental differences across DHI score levels. As Figure 8A shows, those participants with the high DHI scores (above 16) separate well from those with mild scores (0-16). The mild scores are not significantly different (results of two sample t-test between DHI subjects with zero and DHI subject with low score; p-value for NSR *p=.568*, for right-to-left force *p=.657*, for left-to-right *p=.4 > .05*), while the high DHI scatter establishes a reference region to help assess other disorders in relation to the clinical symptoms of this clinical phenotype.

**Figure 8:**
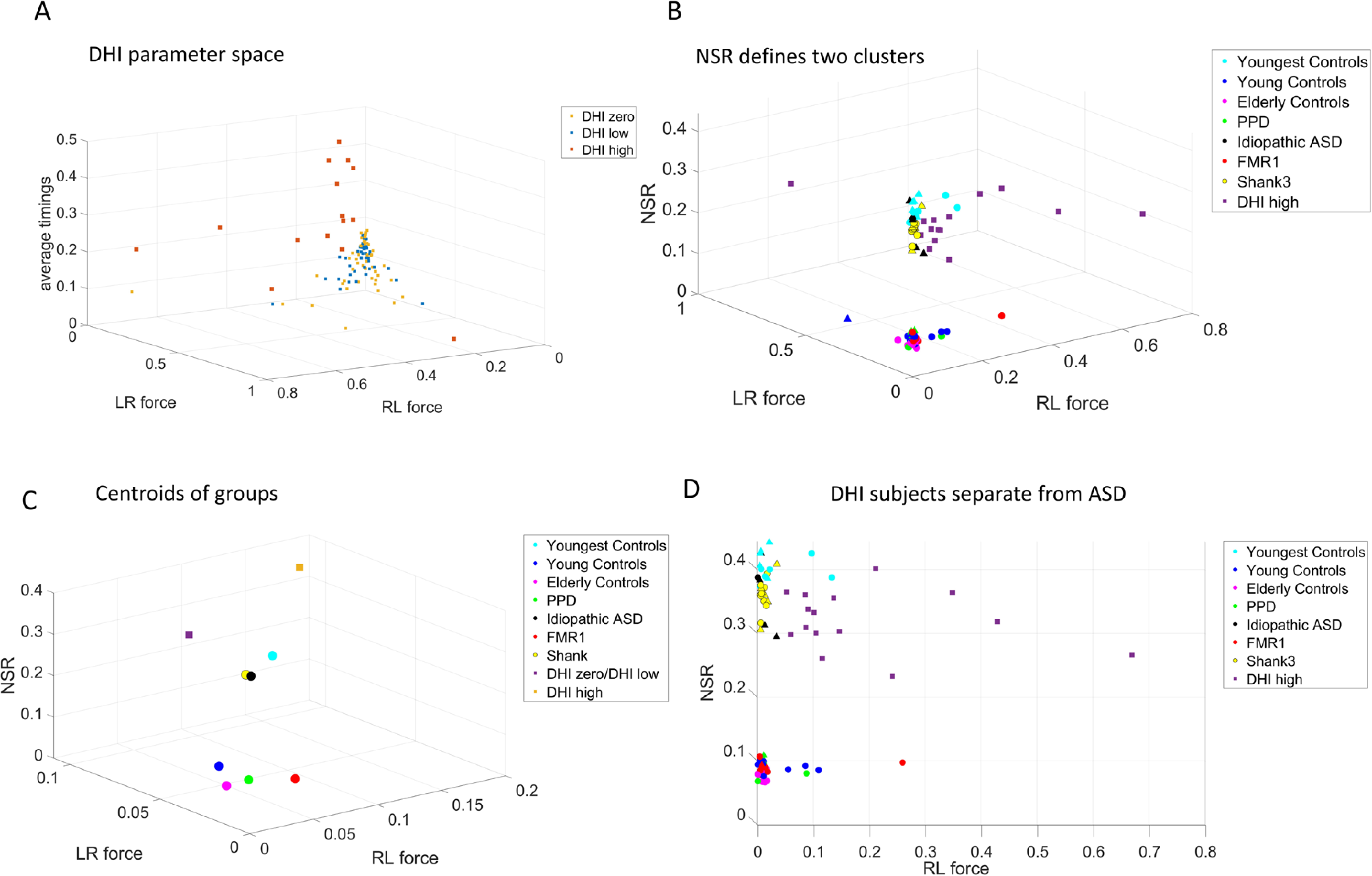
Vestibular DHI vs. subcortical PPD and FMR1 automatically segregate and point at gait differences in idiopathic- and SHANK3-ASD. **A)** Severe vestibular DHI scores above 16 are clinically significant and separate from low-to-mild cases in the parameter space of average timings, left-to-right, right-to-left feet acceleration MMS forces (causality divided by corresponding lags.) B) High vestibular DHI, idiopathic ASD, SHANK3-ASD participants and their youngest age- and sex-matching controls separate along both the NSR and the force axes, from the older controls, the PPD and the FRM1 carriers. In non-DHI participants, circular markers represent females and triangular markers represent males. Two sample t-tests showed no significant differences between males and females. (p-values across x-y-z dimensions for the upper cluster*; .25, .94, .83 > .05*, p-values across x-y-z dimensions for the lower cluster; *.26, .12, .06 > .05*) C) The averaged point (centroid) of each group emphasizes the separation in (B). D) A 2-D projection of the 3-D plot of (B) with all DHI included, shows that despite similarities in the NSR of the vestibular DHI and ASDs, those participants with a high DHI score depart from still-developing youngest participants and from ASD participants (both idiopathic and of known etiology.)

Second, using the high DHI scatter as reference, we define a parameter space spanned by the Noise-to-Signal ratio (NSR) and the causal forces. There, the vestibular DHI participants departed from adult and elderly controls, grouping instead, along the NSR axis, with the youngest, still-developing controls. Their NSR levels matched those of the ASD-SHANK3 deletion syndrome and five Idiopathic-ASD participants that we included as reference (owing to their known high MMS NSR ^32,34.^) The subcortically driven neurodegenerative PPD and the FMR1 carriers separated from the vestibular DHI adults, closer in age than the ASD participants. As such, biomechanical gait noise derived from linear acceleration MMS, automatically separated these gait disorders with different clinical diagnosis (Figure 8B). Figure 8C shows the summary tendencies of each group (obtained by calculating the centroid of each cluster.) Furthermore, we appreciate on the projection plane spanned by the NSR and the right leg (RL) force parameter, that the spread of participants with high vestibular DHI scores is much higher than that of ASD participants and age-matched controls (Figure 8D.) The scatter along the right-to-left causality-to-timings ratio distinguishes SHANK3-ASD, idiopathic-ASD and vestibular DHI disorders, despite the common levels of NSR.

Perhaps this separation could be attributed to the intrinsic difference in neuromotor dynamics between these groups. It seems that while NSR reveals differences and similarities in the phenotypical motor component of a class of heterogenous groups and disorders, causal analysis reveals important underpinning differences in neuromotor dynamics. Finally, even though both SHANK3 deletion syndrome and FMR1 premutation carrier syndrome may often receive a diagnosis of ASD at a young age, they are clearly differentiated in this parameter space.

It is possible that our MMS derived from acceleration paired with these new analytics, distinguish important nuance behaviors currently missed by the naked eye of an observer. The results from these acceleration-based analyses yield support for a separation between disorders that are fundamentally driven by subcortical structures (*e.g.,* PPD, FMR1) and those which might be primarily tied to cortical and vestibular structures (*e.g.,* idiopathic- and SHANK3-ASD). While the former may be formulated within a framework of biomechanical full-body motor control, postural coordination and motor synergies, the latter also include an element of sensory integration (auditory, visual, and proprioceptive signals) and perceptual issues tied to peripersonal space, temporal dynamics, and gravity-specific control.

To examine temporal information within the efference-reafference framework, we calculated for each undirected pair of angular speed MMS time series (*i.e.*, the full time series, not only the peaks) the coupling frequency for which the cross-coherence is maximized. To that end, we found the average of all cross-coherence outputs of a node’s MMS series with all the other MMS series nodes in the network (see Supplementary Materials). The time between two MMS is essentially a form of inter-spike interval timing for the information flow crossing each body node. Therefore, we average for each group the inter-spike times of each node and compare the rate of putative afferent (incoming) and efferent (outgoing) activity. Figure 9 shows the patterns from a series of t-tests between the timings, using every possible combination of the two groups for each node. Zero entries mean that the t-test was not significant, non-zero entries show the average differences between the mean times for each group.

**Figure 9:**
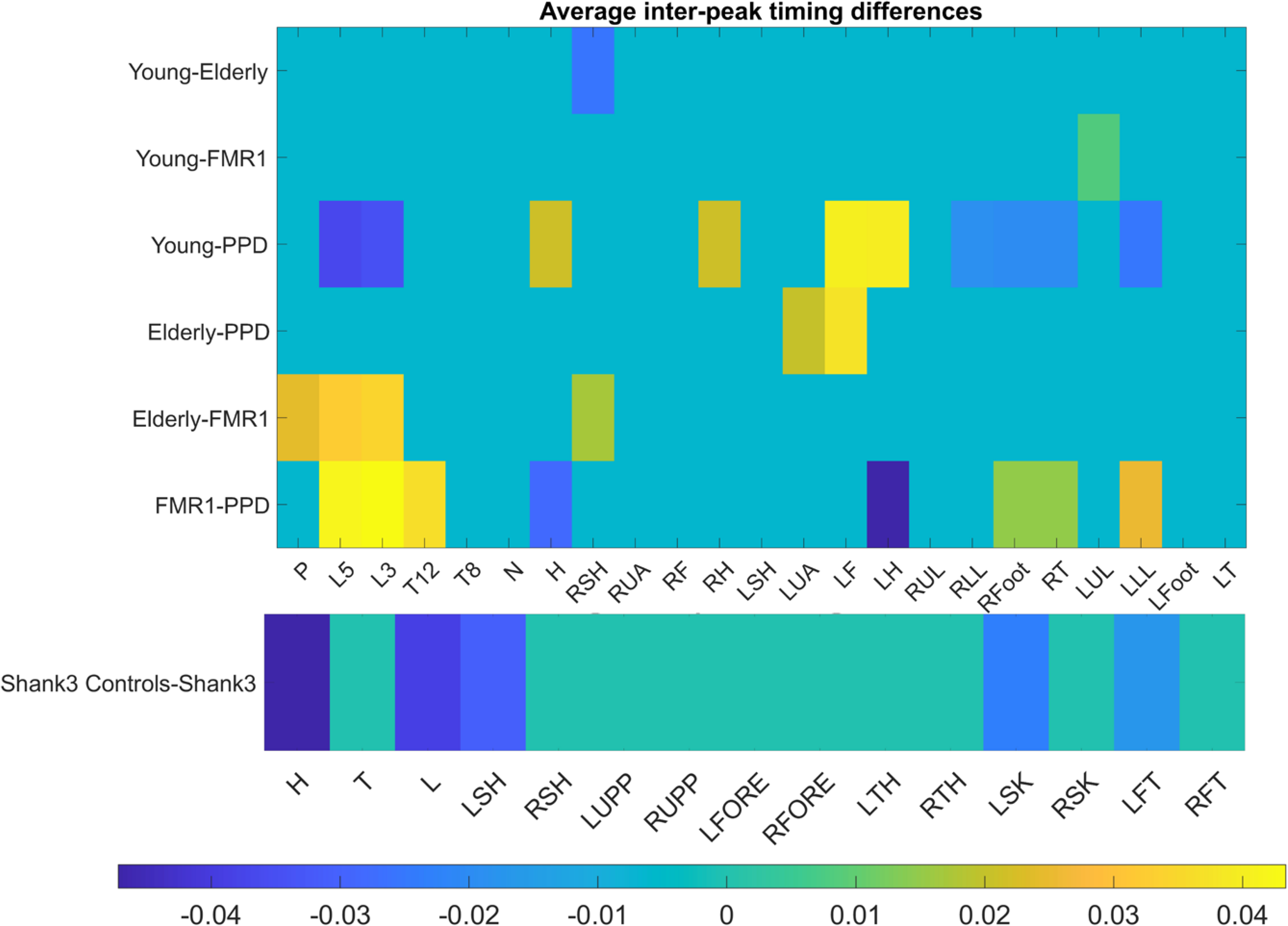
The average differences in inter-peak timings (seconds) taken for each group across all nodes, reveal separation between groups. A difference of zero (cyan color) indicates no significant difference (*p=0.05*).

We note that FMR1 carriers exhibit much higher inter-peak timings in the thoracic area and the lower leg area than the PPD subjects. Both young and elderly controls seem to have much higher inter-peak timings in the (non-dominant) left arm and hand than PPD patients. Interestingly, FMR1 carriers seem to depart further away in their timing profiles from elderly controls than elderly do from the young controls. This suggests that the healthy aging process may not significantly alter the afferent or efferent rate of activity in any specific area of the body, despite increasing the inter-peak interval timing (on average) for the entire body. As we saw earlier (Figure 1) these consistent patterns perhaps hint that aging may not be anatomically specific with regards to the rate of activity. On the other hand, aging seems to decrease the efferent and afferent activity as well as increase the internal motor timings of the system. Further analyses of the SHANK3 cohort (accessing 15 rather than 23 nodes) indicated that for most body parts, young developing controls seem to have higher inter-peak timings than those with SHANK3 deletion syndrome. As we saw in Figure1C, on average, controls have higher inter-peak timings. This is the case here across most nodes, though, for certain body parts, the opposite trend (negative values) can be observed. Most notably, the head node is the most negative, indicating higher inter-peak timings in SHANK3 deletion syndrome (Figure 9.) This is not surprising, given that SHANK3 codes for a scaffold protein located at the post-synaptic density (PSD) of glutamatergic synapses. These are important for the formation and stabilization of synapses, as they assemble glutamate receptors with their intracellular signaling apparatus and cytoskeleton at the PSD ^45^. Thus, delays in signal transmission are expected to be reflected, as we use inter-peak timings as a proxy of this internal process.

Given the differentiation between adult controls and very young participants (including those with SHANK3- and idiopathic-ASD), we then focus our next analyses on the young controls of college age, the elderly controls and the PPD and FMR1 participants, and separately perform the same analyses on the youngest groups of controls and ASDs. The cohort corresponding to the lower NSR cluster in Figure 8B is presented in Figure 10, while the cohort with higher NSR in Figure 8B is presented in Figure 11.

**Figure 10:**
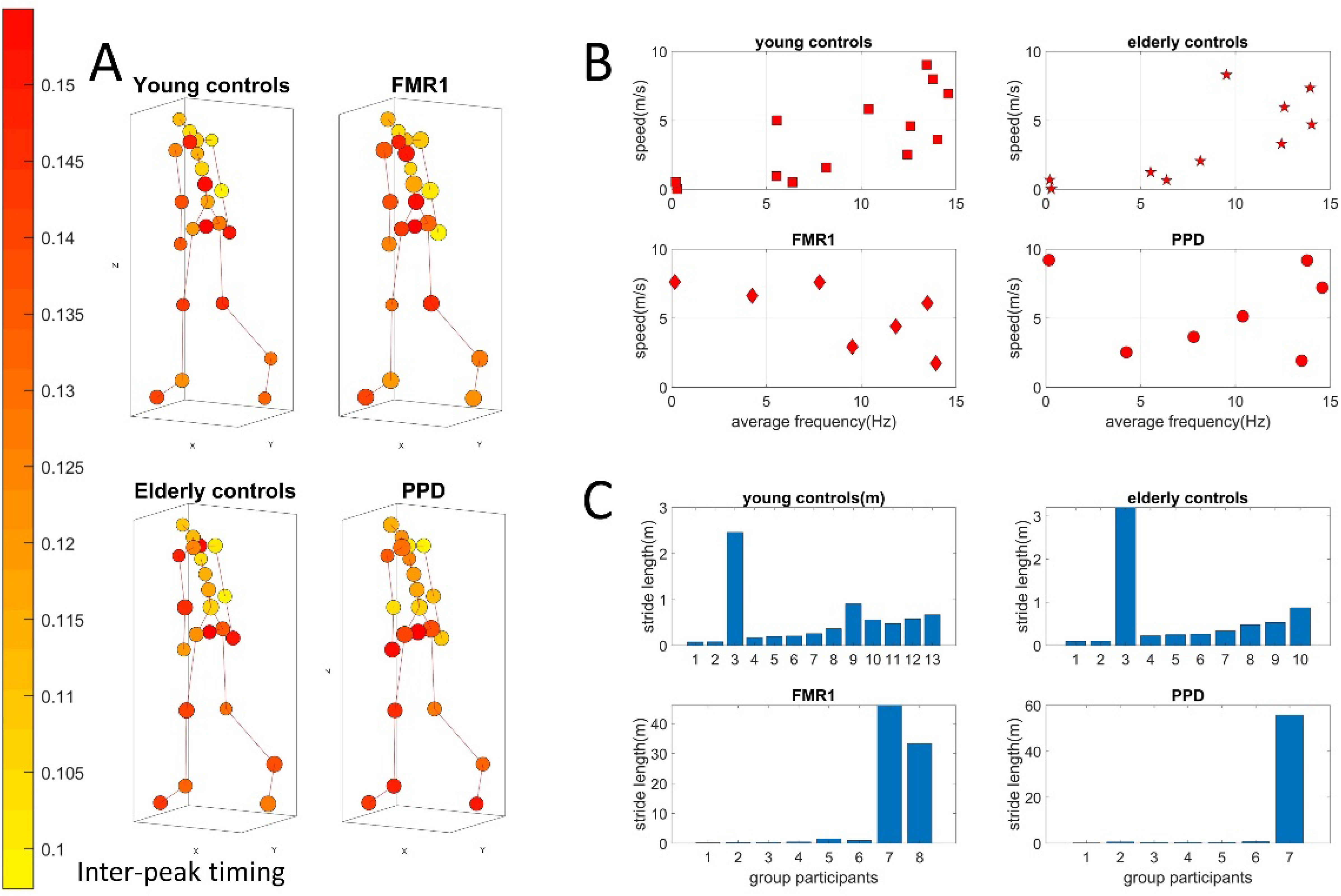
Stride-length differences from speed to frequency ratio obtained from the center of mass trajectories, separate healthy aging from nervous systems disorders. (A) Network representation in anthropomorphic avatar form. The size of the nodes is proportional to the maximum coupling frequency (the one that maximizes cross-coherence) of a node with the rest of the body and the color of the nodes is the average time distance (s) between two MMS peaks. (B) Parameter space comparing speed vs. frequency. Focusing on the trajectory of the center of mass of the lower body, mean speed over mean frequency of the speed time series has a positive upward trend for healthy controls. This contrasts to a negative trend in FMR1-carriers and variable trends in PPD. (C) Both types of trends and variations span a range of values across both dimensions with stride length periodically accumulated differing across groups.

**Figure 11:**
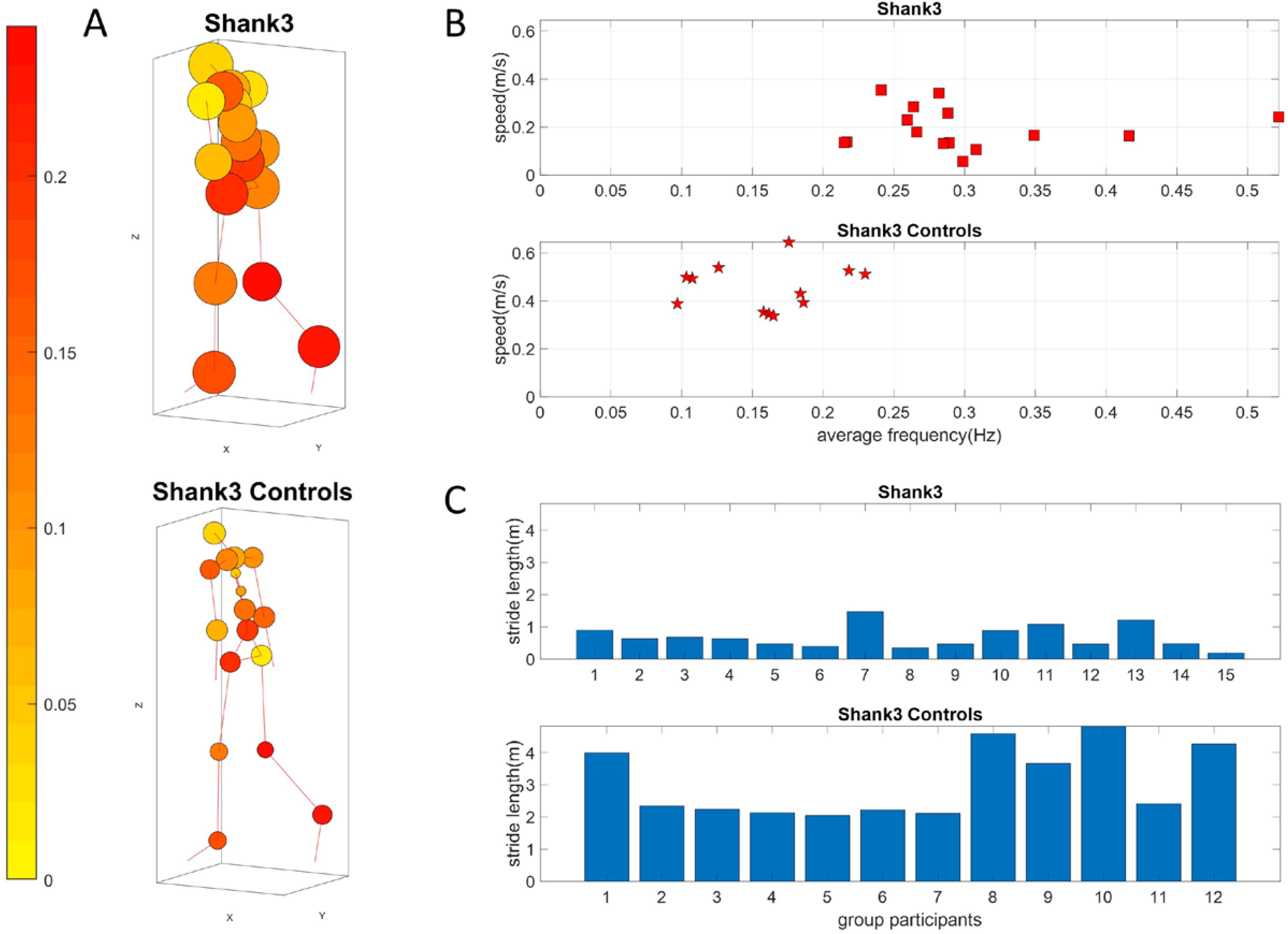
Stride-length differences from speed to frequency ratio obtained from the center of mass trajectories, separate young neurotypical controls from SHANK3-ASD participants. (A) Network representation in anthropomorphic avatar form. The size of the nodes is proportional to the maximum coupling frequency (the one that maximizes cross-coherence) of a node with the rest of the body and the color of the nodes is the average time distance (s) between two MMS peaks. (B) Parameter space comparing speed *vs*. frequency. Focusing on the trajectory of the center of mass of the lower body, mean speed over mean frequency of the speed time series has a positive upward trend for healthy controls. This contrasts to a slight negative trend in SHANK3. (C) Visible and consistent differences in stride length with SHANK3 deletion participants showing an abnormally small stride length in relation to age- and sex-matched controls.

Analyses across the 23 nodes accessible with instrumentation are visualized as an anthropomorphic interconnected network representation in Figure 10A. The figure shows the average inter-peak interval times as the color of the node. The average cross-coherence output of each node’s MMS series is proportional to the size of the node. We appreciate differences between the size of the nodes for the different groups, hence reflecting differences in the maximum coupling frequencies. Methods Figure 13 shows the analytical pipeline to obtain pairwise cross-coherence, along maximal values of the coupling and corresponding phase lags and frequencies employed in the derivation of relevant parameter spaces.

**Figure 12:**
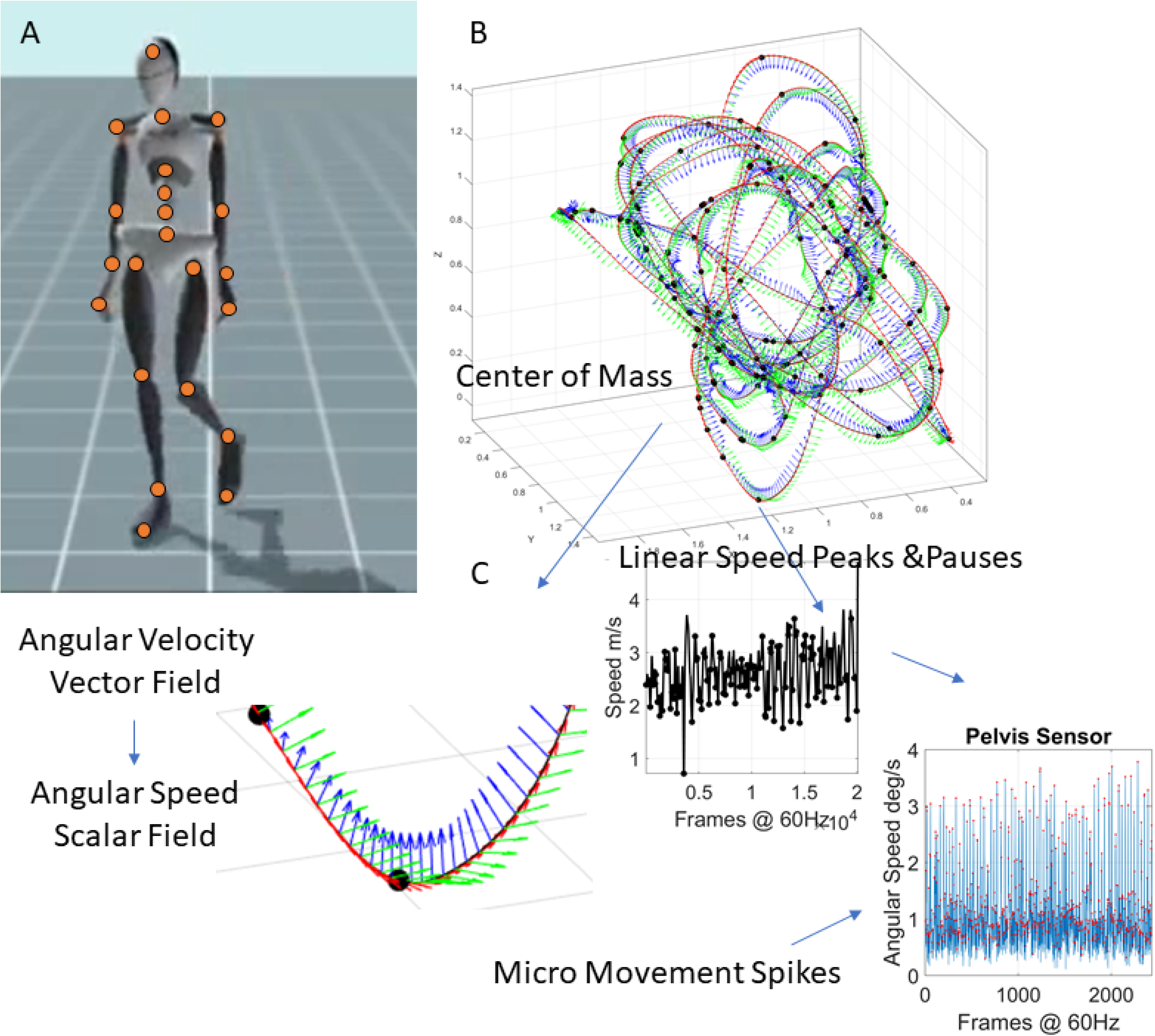
Methods Figure. (A) Data acquisition using a grid of wearable sensors calibrating position and orientation in real time and sampling changes of position and orientation in time, internal sampling rate 1KHz, output 60-120Hz (specs at xsens Mtw Awinda, https://www.xsens.com/products/mtw-awinda). (B) Center of Mass trajectories (m) in 3D parameterized using the Frenet-Serret frame to study geometric aspects of the curve. (C) Linear speed used to mark pauses and peaks along the curve, thus allowing us to express behavioral landmarks along other kinematics parameters such as the angular speed quantifying bodily rates of 23 joints’ rotations. The MMS are derived from the fluctuations in angular speed amplitude. Red dots mark peaks (transitions of speed slope from positive to negative.)

**Figure 13:**
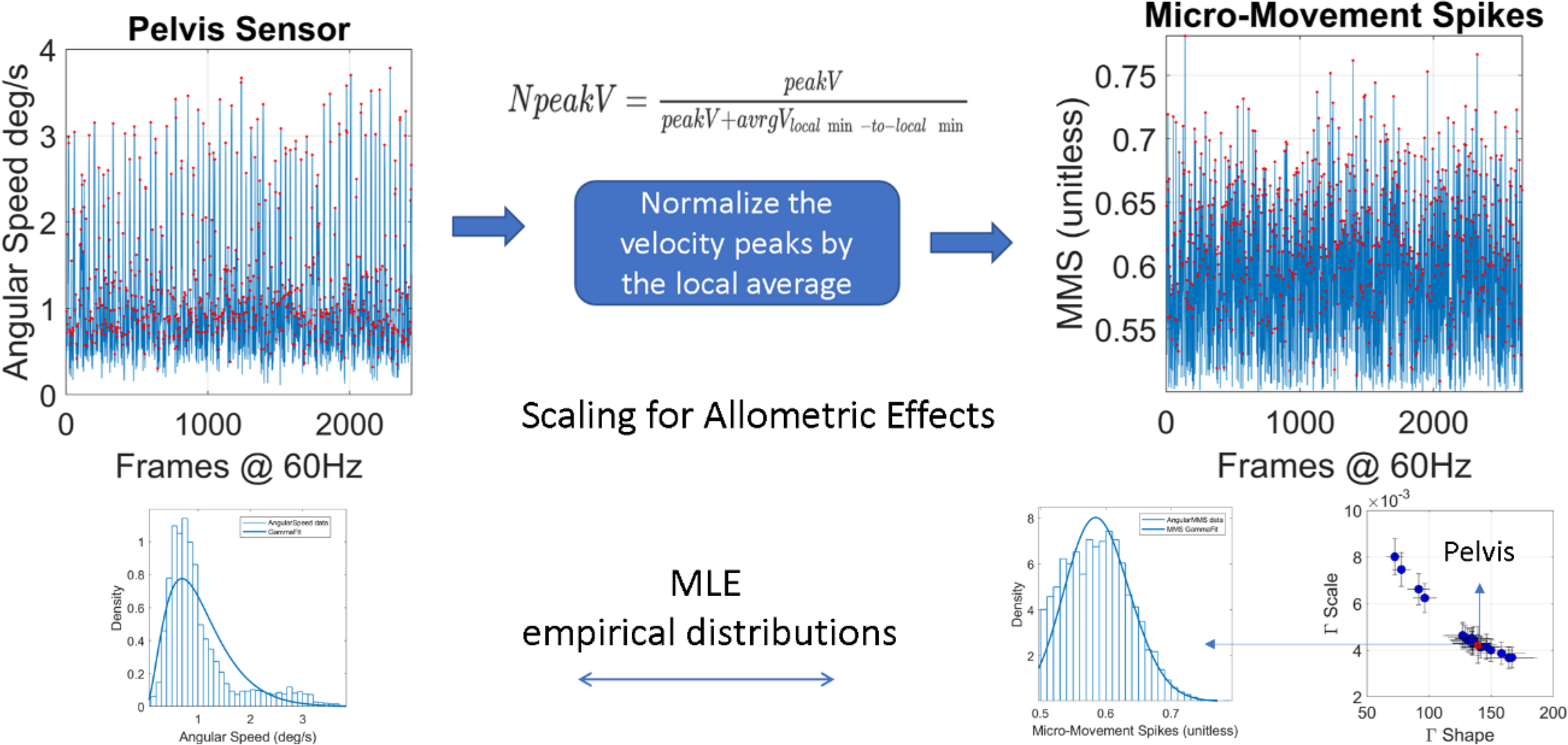
The MMS time series extraction from the angular speed tracking stochastic trajectories from all joints (red dot marks the pelvis for instance.)

For each subject, we calculated the average speed of the COM of the lower body, as well as the average frequency of the speed time series. We did so, to study how the kinematics of a subject relate to the stochastic signatures of the MMS activity. For all groups, except the FMR1 group the speed generally increases with respect to the frequency while for the FMR1 carriers the speed decreases. This is depicted in Figure 10B.

The speed is in m/s and frequency is in Hz, or 1/s. Therefore, speed over frequency has unit of meters. Behind this unit analysis, an important biomechanical concept is “hidden”. Due to the periodic nature of gait, frequency expresses the number of walking cycles or strides per unit time. Speed on the other hand, is the average distance traveled by the center of mass of the lower body per unit time. Distance over time divided by number of walking cycles over time gives us the distance traveled *per* stride. Revisiting Figure 5, we observe that the third dimension, which is speed over frequency does not really differ between participants of each group. This indicates that cumulative distance per stride might be an invariant quantity by which we could characterize each group. Figure 10C shows the different patterns of cumulative distance per stride across participants. We note here that the larger the amount of high frequency jitter in the MMS, the larger the accumulation counting towards the stride length. As such, some participants in the PPD and FMR1 carriers (both of which develop tremor eventually) already show signs of this dysregulation of involuntary jitter in the walking patterns.

## Discussion

This study explored walking activity in the context of natural aging and well-established neurological conditions tied to vestibular and subcortical disruptions. These included patients with a broad range of vestibular Dizziness Handicap (according to the DHI-S inventory) and PPD, FMR1 (Fragile X premutation carriers who may also receive an ASD diagnosis at an early age.) We aimed at examining the patterns of variability from kinematic parameters that would automatically stratify this random draw of the population comprising a multitude of ages and disorders. Furthermore, we explored the movement trajectories of multiple nodes across the body, with the purpose of deriving proxy metrics related to internal temporal lags in information flow. To that end, we framed our problem within the context of causal prediction models of outgoing efferent and incoming reafferent information flow, derived from time series of MMS. We found that using parameter spaces derived from causal connectivity analyses and associated temporal lags, we can stratify a random draw of the population comprising various disorders of the nervous system. The main clusters that automatically self-emerge from our analyses separate vestibular-from subcortical-gait issues and forecast differentiable gait problems in FMR1 participants and the elderly undergoing natural aging.

We distinguish the newly defined internal time lags from external movement times modelled a priori in biomechanics and robotics. Such trajectories’ time profiles are often modelled using the equations of motions to study human movements’ trajectories within some optimization framework (*e.g*., optimal control models such as minimum jerk ^46^, minimum torque rate ^47^, minimum energy ^48^, among others.) Temporal dynamics have been modelled as well in studies of motor control, within the frameworks of stochastic optimal control and reinforcement learning ^49^. In human and animal psychophysics studies associated with reaction times, temporal information is often derived from computer mouse-clicks or lever presses, respectively, within the context of perceptual learning, decision making and other tasks in the cognitive domain ^50^. We also distinguish from those studies the notion of internal temporal lags reported here.

To better appreciate the difference between the internal time lags examined here and those from biomechanics, or from reaction time tasks, we invite the reader to distinguish between a person-centered perspective, *i.e*., obtained in personalized and objective manner, and an observer-centered perspective. The latter makes a statement about the observer’s appreciation of actions’ trajectory timing under some a priori defined population statistics, or grand averaging parametric model. It also imposes a theoretical classical mechanical time-dependency that is incongruent with empirical data. Empirical studies show that the geometry and the forces characterizing motion trajectories are separable. Unlike in inanimate rigid bodies in motion, in biological bodies with nervous systems, the trajectory’s temporal dynamics depend on the level of intent that cortical neurons plan and update online (*e.g.,* ^51^ *vs*. ^38,52–55^). The temporal dynamics of motion trajectories also depend on subcortical regions thought to specify the rate of change in bodily configurations, as the body transitions from one configuration to the next ^56,57,^and we can model it and study it in PPD ^53,55.^In contrast to these centrally defined timings, internal time lags examined here, aim at characterizing temporal dynamics from the efferent and reafferent flow of information across the person’s body, at the periphery. This characterization is done empirically, by estimating the person’s stochastic signatures from the individual fluctuations in the signals’ peak amplitudes and inter-peak-intervals’ timings within the framework of causal prediction in time series forecasting analyses ^33,58.^ Furthermore, our characterization non-trivially augments von Holst’s principle of reafference, to include non-voluntary aspects of actions’ consequences into the feedback that the brain receives from internally self-generated actions.

In the present study, we aimed to study time from a person-centered approach rather than from an external observer approach. This led to self-emerging stratification of the cohort. Instead of a priori assuming a given distribution to characterize the moment-by-moment fluctuations in the walking parameters, we let the data drive our exploration, revealing maximally informative parameter spaces that automatically separated our cohort into clusters that differentiated between temporal dynamics of vestibular and subcortical gait pathologies. While models of biomechanical times cannot account for empirical results in motor control pointing at a fundamental separation between intended and unintended actions, animal and human models of reaction times highlight the subjective nature of time perception during reaction tasks in mammals ^59^. Such models make a general population statement, rather than an individualized characterization of the temporal dynamic’s phenomena.

Indeed, proper parameter spaces allowed us to separate natural aging, PPD and FMR1 premutation carrier syndrome from the baseline of motor activity in young controls. Furthermore, typical, and compromised neurodevelopment (in the form of idiopathic- and SHANK3-ASD) revealed a fundamental departure from gait issues tied to subcortical structures and closer to vestibular dysfunction. Of particular interest was the finding that in natural aging, the lower extremities seem to be compromised, leading to a reduction in the causal prediction of lower body by upper body extremities. This is accompanied by a lengthening of the optimal time lags describing information flow derived from fluctuations in movements. The naturally aging person seems to have communication flow with higher NSR than the younger controls. This result was clearly expressed in lower feedback from the legs and feet to the upper body. The peripheral flow of information appeared to be severely compromised in the elderly PPD and surprisingly so in much younger FMR1 carriers.

The moment-by-moment fluctuations in the peak amplitudes of the MMS derived from linear and angular accelerations offered new ways to stratify this random draw of the population comprising multiple disorders of gait. Of particular interest here is the multimodal nature of the peaks’ distributions revealing a prevalence of the exponential family that increased with natural aging and was much more prominent in FMR1 carriers and PPD participants. In contrast, during early childhood, we see a prevalence of the Gaussian mode that is atypically lower in SHANK3-ASD, manifesting atypically higher exponential mode. This result highlights the random, memoryless nature of the fluctuations in these groups with pathological gait. It also explains the marked reduction in causal prediction of upper-to-lower body and the lack of feedback from lower to upper body found in the FMR1 carriers and in the PPD participants. Importantly, the person-centered approach used in this work was amenable to reveal within each group (developing, young, elderly, PPD, FMR1 carrier, idiopathic- and SHANK3-ASD) different subtypes according to the center of position derived from the lower body kinematics. We used these results to further derive parameter spaces that combined asymmetry in Granger Causality, noise-to-signal ratio, and distribution skewness, to express and to differentiate each group in relation to young controls. Pair-wise differentiation and patterns were also revealed by the inter-peak timing analyses of the trunk, arms, and legs regions, with different trends in natural aging *vs.* aging with a nervous system pathology.

The results from this study are amenable to build digital screening tools using parameter spaces derived from a simple walking task. Furthermore, the work offers a unifying framework to help predict the early appearance of large departure from normative ranges in young controls, both for normal aging and for young participants who are FMR1 carriers. This is important, given the high penetrance of FX-related syndrome in other neurological conditions across the lifespan. Among these are ASD, FXTAS and PD. The methods described here may offer a new way to detect these gait problems 15-20 years ahead of their clinical onset and as such, could help advance neuroprotective intervention models.

While our analysis uncovered the relationship and similarity between PPD and FMR1 carrier syndrome, it also sheds light to the process of natural aging and how individuals with PPD age differently from neurotypicals. Our results indicate that in elderly controls, there is higher overall causal connectivity in the lower body compared to young controls. It is possible that this is related to the degeneration of motor functions in the lower body which comes naturally with aging. To preserve balance and avoid falls, the nervous system will incur in higher cognitive load. As such, natural walking, a process that is rather automatic in young people becomes cognitively effortful and requires higher concentration as we age. This higher cognitive (central) demand may be reflected in the lower body connectivity at the periphery, and poorer lower-to-upper feedback patterns found in the elderly participants.

The results also reveal fundamental differences between PPD and much younger FMR1 carriers, who nevertheless, also showed a marked reduction in feedback from the lower to the upper extremities. These two subgroups were far apart in age, yet both revealed large departure from young controls and timing patterns closer to those of the elderly controls.

The PPD cohort, shows not only a significant increase in lower-body connectivity, but also a marked reduction in upper-body connectivity. This pattern suggests that the degeneration of motor control in PPD is not limited to the lower body, but that in fact, it is a disease that affects the motor system in a general and systemic way. A higher cognitive load is required to control the lower body which as a side effect may result in poor connectivity in the upper body, and poor systemic coordination with impaired kinesthetic sensory feedback. Interestingly, when comparing how the inter-peak timings of each body node differ between groups, we can conclude that in that sense, the elderly controls and the FMR1 carriers are almost identical to young controls. On the other hand, PPDs have significant different inter-peak intervals’ timings than young controls. This not only shows the departure of PPD from natural aging, but also shows that despite the Parkinsonian symptoms of FMR1 syndrome, the latter is still a fundamentally different condition than PD.

To further explore differences in gait pathologies, we examined a large cohort of participants with a broad range of scores from the vestibular Dizziness Handicap Inventory screening (DHI-S). This questionnaire quantifies the person’s overall malaise vertigo, fatigue, and lack of balance. Then, we further characterized these scores using digital data from walking activities, to assess their gait. We used linear acceleration (as it relates to vestibular function) and derived the MMS and similar standardized ratios as with the other participants. Using the feet data, we localize the cohort on the parameter spaces that we designed and automatically separated the high-score (severe) DHI participants from those with lower (low to mild) condition. Using the severe cohort as reference, we localized PPD, FMR1, young and elderly controls along with still-growing young children with typical and atypical neurodevelopment. The latter included idiopathic- and SHANK3-ASD. They grouped with the severe vestibular DHI, along the NSR axis of our parameter space, rather than with the PPD and FMR1 participants and age-matched controls. This result distinguishes vestibular from subcortical gait disturbances and may help us further refine the heterogeneous ASD.

### Future Paths of Inquiry

Finally, as a potential actionable application of the present work to mitigate the high levels of noise found in the walking patterns of ASDs, PPD and FMR1 carriers, we used the relationship that Granger’s causal prediction methods offer between frequency and time domains. Inspired by Granger’s approach, we developed a network-connectivity based model of noise cancellation (Supplementary Figure 1). Specifically, we systematically removed frequencies reportedly associated with tremor in the literature ^60^ and quantified the outcomes of this systematic removal on the noise portrait derived from the time series analyses of angular speed peaks of walking patterns. We found differing ranges of optimal tremor removal relative to healthy young *vs.* elderly controls group. The former revealed optimal ranges for frequencies above 19 Hz, while the latter showed frequency band with lower limits, near 10 Hz. More exploration of these aspects of noise cancellation are warranted, yet using the present approach exploring internal optimal time lags within the context of causal prediction, we open new avenues of personalized support tailored to differentiable gait issues in neurodevelopment (ASD) and neurodegeneration (PPD) with forecasted problems in FMR1 carriers.

Besides the frequency-time relations highlighted in the above exploration, the work also revealed important spatio-temporal features of the MMS. Namely, through examination of the joint empirical distributions of MMS derived from fluctuations in the angular speed peaks’ amplitude and the MMS derived from the fluctuations in inter-peaks-interval timings, we found interdependences and complex patterns rendering invalid the common assumption of independence in the spikes data (Supplementary Figure 2). This result also opens new questions about common assumptions of stationarity and homogeneity in the data. Future empirical and theoretical work will be necessary to advance this important area of research, vital for interpretation and statistical inference of behavioral phenomena.

### Conclusions

In summary, we offer a new unifying framework to stratify a random draw of the population based on natural walking patterns, identify subtypes withing each group and differentially anticipate gait pathologies of the nervous systems bound to appear in neurodevelopment, natural aging and emerge from genetic-based constraints. Furthermore, our simple models are computationally viable for real time implementation. As such, we suggest actionable models of noise cancellation to build supportive interventions tailored to different subgroups of vestibular and subcortical gait pathologies. Our work warrants investigation on computational models to help enhance causal prediction and feedback from movements and to improve internal temporal lags of information flow.

## Materials and Methods

This study was approved by the Rutgers and the Columbia University Institutional Review Boards (IRBs) abiding by the precepts of the Helsinki Act. A total of 187 participants were included in the analyses (See Supplementary Table 1 for general demographics and Supplementary Tables 1-5 for demographics and clinical information in each group.) Patients with Parkinson’s Disease (PPD) were recruited through the clinical_trials.gov site, the Robert Wood Johnson clinic. Participants with Dizziness Handicap were recruited through Columbia University, during their visit to the Otology and Neurotology clinic. ASD participants were recruited from the New Jersey Autism Center of Excellence (https://www.njace.us/) and SHANK3 deletion ASD participants were part of a clinical trial in collaboration with Mount Sinai. Informed consent was obtained from the participants according to our Rutgers and Columbia University IRB-approved protocols. Different aspects of these data unrelated to stochastic gait analyses, were previously published ^7,28,61,62.^

### 2.1 Experimental Design and Statistical Analyses

Participants (PPD, ASD, FMR1 and Controls of all ages) wore the XSens system (17 wearable sensors across the body collecting position, orientation, and acceleration at an internal sampling rate of 1000Hz with an output rate of 60-100Hz, and the SHANK3 deletion syndrome participants wore 15 wearable sensors, Polhemus Liberty output sampling at 240Hz, co-registering data streams from 15-23 joints across the body, depicted in Figure 12A). In-house MATLAB routines were used to implement the Frenet-Serret frame ^63^ and derive various kinematics parameters. Position was used to derive linear velocity and linear acceleration fields, which were converted to scalar fields of speed (m/s) and acceleration (m/s^2^) respectively. Then, maxima and minima were obtained as landmarks to track the speed of the center of mass (e.g., Figure 12B) (obtained by replacing mass with length in the original equation 1).

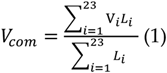

Participants in the vestibular DHI cohort wore SoleSound, a fully portable research prototype developed at the Columbia University Robotics and Rehabilitation Laboratory and previously validated as an instrumented footwear system for the measurement of spatiotemporal gait parameters ^28^. There are two nine-DOF inertial measurement units in each footwear unit: one placed inside the sole of the sandal along the midline of the foot below the tarsometatarsal junctions, and the second is in a small plastic housing which is then secured with a Velcro strap to the anterior aspect of the subject’s lower leg, below the knee. Four piezo-resistive force sensors are additionally located on the sole of each footwear unit to measure the pressure beneath the calcaneus, the head of the fifth metatarsal, the head of the first metatarsal, and the distal phalanx of the hallux. Together, the piezoresistive sensors and the inertial measurement units measure pressure under the foot and kinematic data of the foot and shank, including the ankle plantar-dorsiflexion angle. An ultrasonic unit is mounted on the posteromedial side of the sole and enables the system to estimate the base of walking. The hip pack unit contains a single-board computer that synchronizes and processes raw data at a sample rate of 500 Hz. We used the kinematics data (acceleration) in our analyses.

Time series of linear and angular accelerations were used to derive the standardized micro-movement spikes (MMS), to serve as input to further analytical pipelines (Figure 12C). Then various ratios and standardized indexes were obtained, to allow for examination of full body and feet behaviors as available, across the full cohort of 187 participants, independent of their anatomical differences and sensor types.

Micromovement Spikes (MMS): The MMS are defined as the normalized peaks of any kinematics time series of the fluctuations in amplitude, as the signal deviates from the mean amplitude. Specifically, to obtain the micromovements time series from the angular speed time series we first calculate the peaks. Each peak is then normalized by the local average speed of the speed, *i.e.,* the average value of the two minima before and after the peak to scale out anatomical differences owing to disparate age and physical dimensions (equation 2).

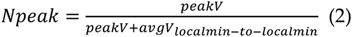

It has been shown that the resulting MMS time series follows the generalized gamma distribution in human motion and applied to gait ^34,64.^Then, from the resulting time series we subtract the gamma fitted mean (fitted using maximum likelihood estimation, MLE). When tracking the MMS continuously, we can used them as a peripheral proxy of the efferent an afferent activity of the central nervous system. Figure 13 shows the pipeline describing the MMS extraction from the sensor data.

#### Cross-coherence

The cross-coherence between two times series (assumed to be the realizations of unknown stochastic processes) is defined as the cross-spectral density between the two series normalized by the product of their auto-spectral densities ^<u>65</u>^. Since human motion is non-linear, in this study we use cross-coherence to quantify the similarity between any two body nodes’ MMS time series in the frequency domain. We then identify the frequency for which the cross-coherence function is maximized. Figure 14 shows the analytical pipeline for the calculation of maximal cross-coherence.

**Figure 14:**
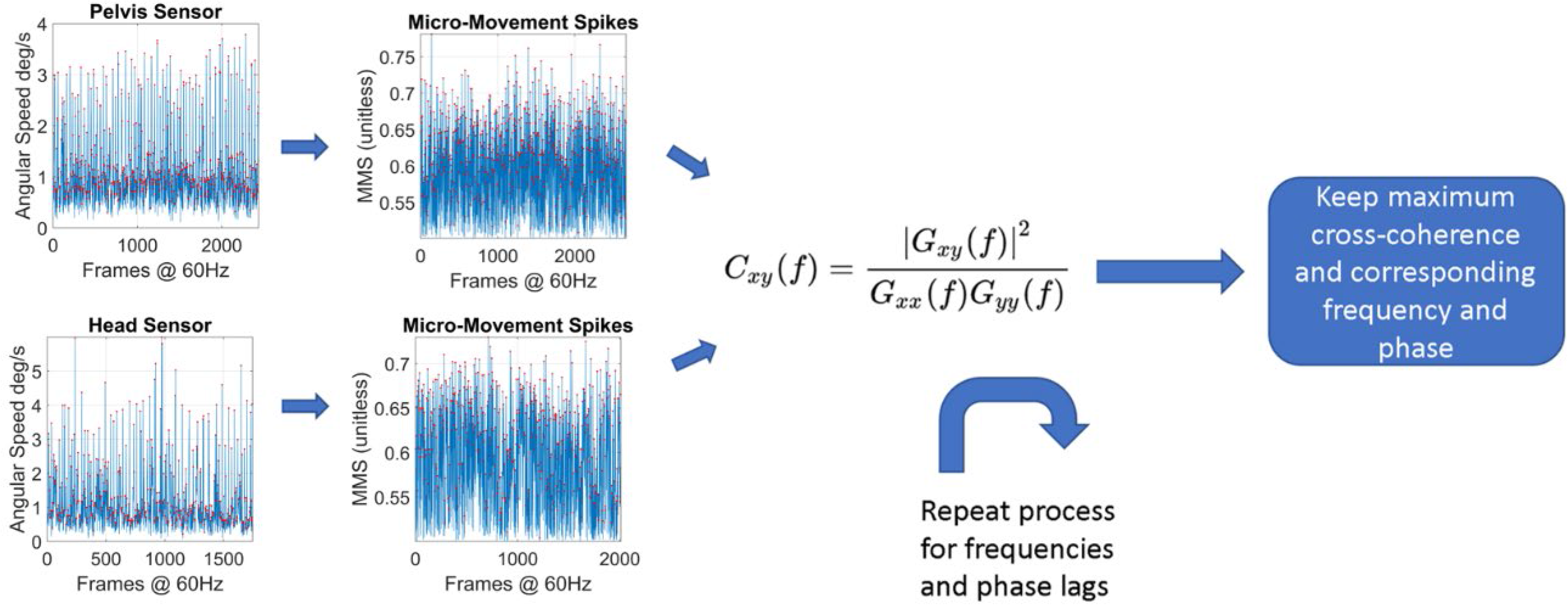
The pipeline for the calculation of the maximum cross-coherence and the frequency for which it is maximized.

**Figure 15:**
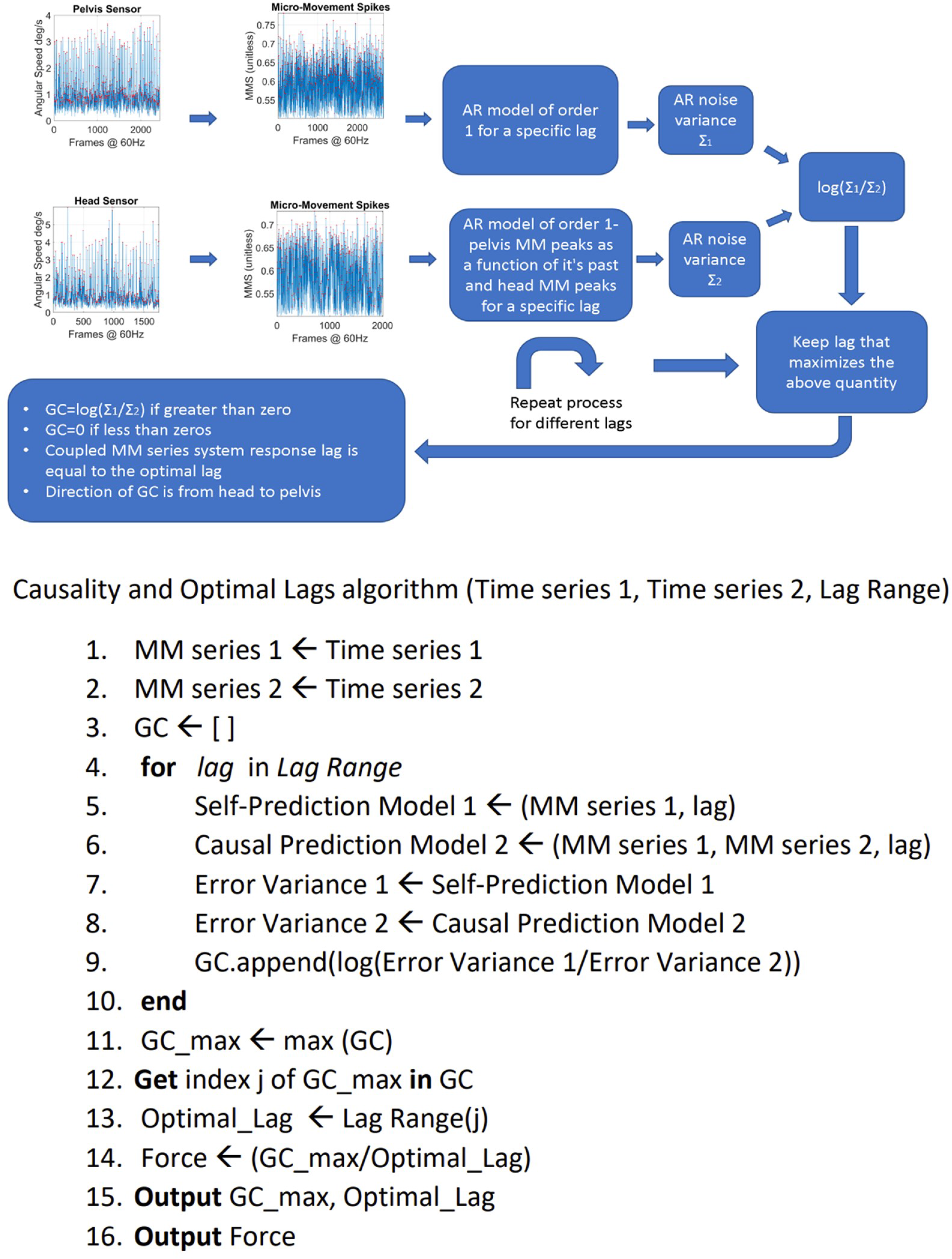
Pipeline for the estimation of the Granger Causality, Response time values and Force between two body parts.

#### Mean frequency

The mean frequency of a spectrogram *P*(*f*) is calculated as:

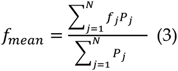

Where *f*_*j*_ is the central frequency of the j-th bin of the spectrum and *P*_*j*_ the corresponding value of the power spectral density. N is the total number of bins ^<u>66</u>^.

#### Granger causality

The MMS serve as a proxy to reflect the inner activity of the nervous system as it is expressed by the kinematics of the various body parts. Hence, we now have a network of times series (23 nodes) that provide information about the mechanisms of the CNS to control the peripheral activity for each subject and each group of subjects. Specifically, we are interested in the relationship between two time series of MMS. Traditional concepts, such as mutual information, cross-correlation and cross-coherence are being widely used to estimate the mutual information between two stochastic processes ^67^, or to investigate how one relates to the other in the time or frequency domain ^62,68,69.^However, they provide no information about the direction in which the information flows, i.e., which stochastic process is the cause and which the effect.

Other concepts such as Transfer Entropy (TE) and Granger Causality (GC) give us both the direction in which the information flows, as well as the quantity of the information flowing from the causal stochastic process to the effector stochastic process ^70^. In the case of Gaussian processes these two are equivalent ^71^. TE does not require a model and is based on information theory concepts. GC on the other hand, requires fitting an AR model of order N to the time series data.

For the purposes of this paper, we choose the concept of GC. There are two reasons for our choice. First, upon problem simulations, GC proved to have a much smaller computational complexity than TE, to be amenable for real-time implementation of our proposed methods. Second, assuming an autoregressive model can help us estimate the response lags between different body activities. We define the response lag as the number of MMS peaks needed for information to flow from the afferent input of a body part to the efferent output of another body part. We note here that the simple AR model that we chose here is just the beginning of our investigation in humans, but other models of causality have been explored in other problem domains (e.g., see ^72^.) Furthermore, previous work has paved the way to address the complex problem of causal delays ^73^. Here we choose different approaches to adapt these concepts and simplify them to study stochastic and non-linear complex dynamical patterns of human gait. While our goal of seeking automatic stratification of a random draw of the population is more modest than that of causal coupling delay detection, we still face several computational challenges. The present is a first attempt to address the challenge of characterizing a multiplicity of gait-related pathologies across the human lifespan using standardized indexes amenable to localize all participants on a common parameter space and reveal self-emerging clusters. As such, we here explore a simpler digital screening, rather than a digital diagnostics tool.

To explain how we estimate the response lags between the activities of any two body parts we must first explain the concept of GC: Granger, in his original paper defined causality for a pair of stochastic processes X and Y in a solid, mathematical way ^42^. Assume we have a closed system or “universe” in which all available information is contained in the stochastic processes X and Y and let X_t_ and Y_t_ be the realizations of these stochastic processes up to time t. Let U_t_ be all the information accumulated from both time series up to *t-1*. U-Y implies all information apart from Y. Granger gave the following definition of causality. Using his own words ^42^:

“If *σ*^2^(*X*|*U*)<*σ*^2^(*X*|*U* − *Y*), we say that Y is causing X, denoted by Y_t_ => X_t_. We say that Y_t_ is causing X_t_, if we are better able to predict X_t_ using all available information than if the information apart from Y_t_ has been used.”

The operator *A*|*B*, where A, B are stochastic processes means the “construction” of a model where A is predicted from B. The operator *σ*^2^(*A*|*B*) means the variance of the error of the model *A*|*B*. Granger’s definition implies that if a model that predicts the dynamic behavior of X that does not include Y, has a greater on average error than a model that includes Y, then Y “causes” X in that sense. Furthermore, Granger gave a formal definition of “causality lag”, which simply informs us how far in the past we should consider samples of X and Y in our prediction to get an optimal prediction.

“If Y_t_ => X_t_, we define the (integer) causality lag m to be the least value of k such that *σ*^2^(*X*|*U* − *Y*(*k*))<*σ*^2^(*X*|*U* − *Y*(*k* + 1)). Thus, knowing the values Y_t-j_, *j=0,1,…m-1*, will be of no help in improving the prediction of X_t_.”

Simply put, the concept of causality lag refers to the maximum lag in the past of the realization of the stochastic process Y, beyond which, including more past samples will result in a bigger prediction error.

Granger also referred to “instantaneous causality”, which simply means that in the prediction we also include the present values of the stochastic processes. In our present work, due to the non-instantaneous nature of neural information, we assume that prediction should include only the past values. In his solution to the problem of estimating causality for econometric time series, Granger assumed autoregressive models of some appropriate order and used techniques of spectral analysis ^58^. He defined two models, one simple causal model that includes the past samples of both time series and one non-causal model that includes only the effector process. For our purposes, we formulated our own model and approach to Granger causality which allows us to quantify the internal motor timings of the system.

Granger Causality has been used extensively in many applications over the years. In a recent work, a simplified Granger Causality map was used to identify root causes of process disturbances. A directed causality graph was built using autoregressive models between operating units and the weights were adjusted based on F-values of parametric granger causality F-tests and then a maximum spanning tree algorithm was implemented to detect causes of process disturbance propagation across the units. This shows that Granger Causality can be employed on a network level where there are multiple processes that interact with each other, such as in our case ^74^. For our purposes, we formulated our own model and approach to Granger causality which allows us to quantify the internal motor timings of the system. We decided not to perform parametric tests and avoided using high order autoregressive models precisely because we wanted to develop a simpler technique that could run on real time and have a direct biological interpretation with regards to the motor timings of the CNS. Clearly, this is a first attempt to model these phenomena and as such, further attempts to continue our investigation on automatic stratification and screening are warranted.

Let X and Y be the two univariate stochastic processes and X_n_, Y_n_ the corresponding time series we obtained from our sensors. For a choice l of the internal time lag of the system, we assume the non-causal and causal models:

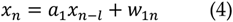

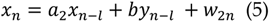

Where *w*_1_, *w*_2_ are assumed to be independent discrete time AWGN (additive white gaussian noise) processes and *a*_1_, *a*_2_, *b* constants we want to estimate from the statistical properties of our data. We also make the hypothesis that the noise processes are independent from X and Y. Assuming that the noise processes are gaussian implies they have zero means. To apply this model without loss of generality we simply pre-process our data by centering them around zero. Note that we assumed in our model independence of the processes, but it would suffice to assume them uncorrelated, which is weaker condition than independence.

To proceed, we multiply equation (4) with *x*_*n*_ and we get:

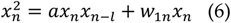

Applying the expectation operator *E*( .) on both sides of equation (6):

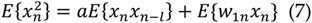

Since noise *w*_1_ and the process X are independent, it means they are uncorrelated. Therefore:

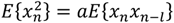

From which we get an estimate for the constant *a*:

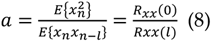

Where *R*_*xx*_(*τ*) = *E*{*x*_*n*_*x*_*n−τ*_} is the autocorrelation function of X.

We continue by multiplying (5) with *x*_*n*_:

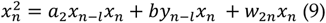

Also, we multiply (5) with *x*_*n*−1_:

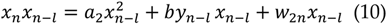

Taking the expectation on both sides of (9) and (10), since X and Y are uncorrelated, we finally arrive to the system of equations:

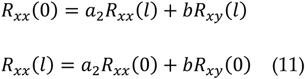

Where *R*_*xy*_(*τ*) = *E*{*x*_*n*_*y*_*n*−*τ*_} is the cross-correlation function between X and Y.

Assuming (11) has a solution, we can estimate *a*_2_, *b*.

The final step in our analysis is to find the variances of the noises.

We square equations (4) and (5):

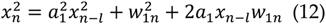

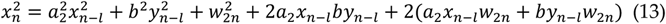

Applying the expectation in both sides, since the noises are assumed to be independent of the processes, all cross-correlation terms between processes and noises are zero, therefore:

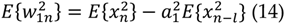

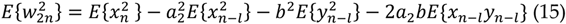

Since the variance of a noise processes *w* is *σ*^2^ = *E*{(*w* − *w*_*mean*_)^2^}, in our case where the noises are zero mean, eventually:

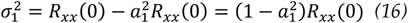

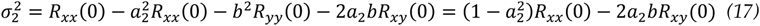

Thus, solving system (9) gives us the constants of the system for a given choice of lag and then equations (16), (17) give us the errors of the non-causal and causal models. Following the logarithmic definition of causality ^75^:

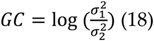

If the causal model better predicts the behavior of X for a given internal motor lag 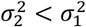 then *GC* > 0.

A standard causal autoregressive (AR) model of order one is of the form:

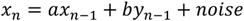

Which is order 1. However, what is the physical meaning of “order 1”? It means we assume the previous sample, one lag in the past of X, contribute to the prediction of X. It is an arbitrary quantity when we consider it independently of the sampling rate *F*_*s*_ of our sensors. In context however, one lag is 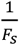 seconds, which is the fundamental unit of time of our system. In our model, we assume:

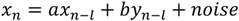

This can still be assumed an autoregressive model of order 1 in which the fundamental unit of time is not one sampling period but *l* sampling periods. Different choices of *l* will give autoregressive models for different ***time scales.*** The time scale *l* for which causality is maximized is our definition of “internal motor timing”, because at this time scale, we have the maximum flow of information from Y to X.

We make the following hypothesis. At any given moment, the process Y_t_ transmits information to the process X_t_ through the CNS. The response lag between the two processes is the time needed for information to travel from the afferent process Y_t_ to the efferent process X_t_. Following this train of thought we can assume that the lag parameter of the AR models that maximizes the GC is the best estimation of the corresponding response time between X_t_ and Y_t_. Therefore, we propose the following pipeline of analysis to find the maximum GC and estimate the response lags between any two sensor MMS time series of the subjects (Figure 14).

Finally, we define “force” as the ratio between maximum GC and the corresponding optimal lag. We propose that this quantity is indicative of the speed of information transmittance, as it considers both the causal influence of one time series on another time series as well as the rate at which it takes place.

## Author Contributions

T.B. and E.B.T. conceptualized the study, designed the methodology and wrote the software. T.B. did the equations derivation, validation, investigation, and formal analysis. T.B and E.B.T. designed the figures / visualization tools. R.R. and J.R. did the data collection and curation of PPD, ASD and Controls. DZ, SKA and AKL provided the DHI data set and did all the data collection and clinical evaluations of the cohort. T.B. and E.B.T. did the original writing— original draft preparation, and all authors reviewed and edited the MS to its final state. E.B.T. did the project supervision, administration, and funding acquisition. All authors have read and agreed to the published version of the manuscript.

## Funding

This research was funded by The New Jersey Governor’s Council for the Medical Research and Treatments of Autism, grant number CAUT15APL038 and by the Nancy Lurie Marks Family Foundation Career Development Award to EBT.

## Acknowledgments

We thank all participants in the study.

## Conflicts of Interest

The authors declare no conflict of interest. The funders had no role in the design of the study; in the collection, analyses, or interpretation of data; in the writing of the manuscript, or in the decision to publish the results.

## Notes

### Competing Interest Statement

The authors have declared no competing interest.

### Summary of Updates

This is a major revision of the original MS. We have added participants of other disorders involving gait. These are 118 with vestibular dizziness handicap, 16 with SHANK3-deletion syndrome and autism, 5 with idiopathic autism, 10 very young controls 6-13 years old and a Tourette's autism participants with visible motor ticks interfering with his gait. We have added several new figures unveiling parameter spaces that stratify this cohort of 187 people into fundamentally different gait disorder types (vestibular-cortical vs. subcortical degeneration) and provided evidence for a new type of temporal dynamics linked to causality optimal lags. Overall the results suggest a new screening tool for gait derived from a simple walking task that can also forecast issue in the horizon of typical aging and aging with FMR1 premutation (in carriers of the FX gene.)

