## Supplementary Material for "Optimal Time Lags from Causal Prediction Model Help Stratify and Forecast Nervous System Pathology"

**Supplementary Table 1: Summary Demographics (All groups)**

| | Mean Age | Females (Total) | Males (Total) |
|---|---|---|---|
| **Young Controls** | 10 (+/- 4) | 4 | 4 |
| **College Aged Controls** | 19 (+/- 2) | 13 | 2 |
| **Adult Controls** | 29 (+/- 4) | 6 | 2 |
| **Elderly Controls** | 58 (+/- 13) | 3 | 1 |
| **Tourette's ASD** | 8 | - | 1 |
| **Idiopathic ASD** | 8 (+/- 3) | 1 | 3 |
| **Fragile X Carriers** | 37 (+/- 21) | 6 | 1 |
| **DHI Participants** | 73 (+/- 8) | 61 | 57 |
| **SHANK3 Participants** | 8 (+/- 3) | 8 | 7 |
| **Parkinsonian Participants** | 69 (+/- 5) | 3 | 4 |
| **Total (187)** | | 105 | 82 |

**Supplementary Table 2: Summary Demographics DHI Group**

| | N=118 |
|---|---|
| **Age mean (SD)** | 73.25 (8.38) |
| **Sex n (%)** | |
| Male | 57 (48.3) |
| Female | 61 (51.7) |
| **Race/Ethn n (%)** | |
| NH-White | 66 (55.9) |
| Others | 52 (44.1) |
| **DHI-S mean (SD)** | 6.29 (8.77) |
| Mild 0-12 n(%) | 96 (81.4) |
| Moderate 14-24 n (%) | 15 (12.7) |
| Severe 26-40 n (%) | 7 (5.9) |
| **Chief Complaints** | |
| Hearing loss n (%) | 99 (83.9) |
| Tinnitus n (%) | 48 (40.7) |
| Dizziness n (%) | 53 (44.9) |
| Others n (%) | 19 (16.1) |

DHI-S indicates dizziness handicap inventory—screening version; NH-White, non-Hispanic white; Race/Ethn, race/ethnicity; SD, standard deviation.

**Supplementary Table 3: SHANK3-ASD demographic and clinical information**

| Subject | Age yr:mo | The Mullen Scales of Early Learning [1] | | | Vineland Adaptive Behavior Scales-II | | | | | ADI-R [2] | | | ADOS-2 [3] | | DSM-IV/ DSM-5 | Consensus Diagnosis |
|---|---|---|---|---|---|---|---|---|---|---|---|---|---|---|---|---|
| | | Gross Motor | Fine Motor | NVDQ estimate [4] | Motor | Com munic ation | Sociali zation | Daily Living | ABC | A: Soc | B: Com | C: RRB I | SA | RRB | | |
| 1 | 5:11 | 18 | 14 | 17.79 | 56 | 42 | 51 | 40 | 45 | 19 | 13 | 1 | 18 | 18 | ASD | ASD |
| 2 | 8:5 | 16 | 13 | 17.81 | 54 | 38 | 48 | 48 | 44 | 23 | 14 | 6 | 19 | 19 | ASD | ASD |
| 3 | 6:4 | 27 | 28 | 21.12 | 61 | 44 | 59 | 51 | 52 | 19 | 10 | 2 | 14 | 14 | ASD | ASD |
| 4 | 5:11 | - | 22 | 27.46 | 59 | 49 | 57 | 48 | 51 | 11 | 13 | 3 | 7 | 7 | ASD | ASD |
| 5 | 6:1 | 18 | 9 | 13.69 | 54 | 36 | 48 | 34 | 41 | 28 | 14 | 7 | 17 | 17 | ASD | ASD |
| 6 | 7:10 | 20 | 15 | 15.43 | 49 | 45 | 48 | 51 | 48 | 30 | 14 | 6 | 20 | 20 | ASD | ASD |
| 7 | 7:8 | 33 | 33 | 8.68 | 56 | 45 | 53 | 51 | 50 | 23 | 13 | 6 | 16 | 16 | ASD | ASD |
| 8 | 9:1 | 13 | 15 | 17.09 | 81 | 54 | 66 | 63 | 61 | 5 | 6 | 4 | 15 | 15 | ASD | ASD |
| 9 | 14:8 | 29 | 28 | 19.39 | 56 | 35 | 32 | 35 | 31 | 16 | 12 | 8 | 20 | 20 | ASD | ASD |
| 10 | 5:11 | 22 | 10 | 9.86 | 56 | 36 | 51 | 38 | 43 | 24 | 12 | 5 | 18 | 18 | ASD | ASD |
| 11 | 15:9 | - | 6 | 4.22 | 56 | 33 | 37 | 28 | 29 | 28 | 14 | 2 | 17 | 17 | ASD | ASD |
| 12 | 5:0 | 14 | 6 | 26.67 | 43 | 40 | 51 | 41 | 43 | 25 | 14 | 8 | 4 | 14 | ASD | ASD |
| 13 | 8:7 | 15 | 13 | 12.01 | 43 | 47 | 48 | 48 | 47 | 28 | 14 | 2 | 5 | 10 | ASD | ASD |
| 14 | 5:9 | 22 | 21 | 34.06 | 54 | 61 | 61 | 55 | 55 | 13 | 15 | 0 | 16 | 4 | ASD | ASD |
| 15 | 5:1 | 21 | 17 | 36.07 | 56 | 52 | 51 | 43 | 49 | 23 | 14 | 6 | 20 | 5 | ASD | ASD |
| 16 | 12:6 | 30 | 20 | 13.51 | 64 | 42 | 42 | 50 | 44 | 29 | 13 | 4 | 22 | 3 | ASD | ASD |

[1] The Mullen Scales of Early Learning; Gross Motor and Fine Motor scores provided are the age equivalent in months
[2] ADI-R cutoff scores for autism are: Social = 10, Communication (nonverbal) = 7, Repetitive Behaviors and Restricted Interests = 3
[3] All participants received a Toddler Module or Module 1 of the ADOS-2

Abbreviations: ABC, Adaptive Behavior Composite (Vineland); ADI-R, Autism Diagnostic Interview-Revised; A: Social, Qualitative Abnormalities in Reciprocal Social Interaction; B: Com, Communication; C: Repetitive and Restricted Behaviors; ADOS, Autism Diagnostic Observation Schedule; Com Total, ADOS-2 SA, Social Affect Total, ADOS-2 RRB, Restricted and Repetitive Behavior Total; NVDQ, Nonverbal Developmental Quotient

[4] The Mullen Scales of Early Learning; DVIQ scores were calculated by averaging the age equivalent for fine motor and visual reception subtests then dividing by the child's age in months and multiplying by 100 (Bishop, Guthrie, Coffing, & Lord, 2011)

**Supplementary Table 4: Idiopathic ASD and Age-matched and older controls**

| Subject | Chronological age (years) | Sex | Subject Type |
|---|---|---|---|
| 17 | 12 | M | Idiopathic ASD |
| 18 | 11 | M | Idiopathic ASD |
| 19 | 10 | M | Idiopathic ASD |
| 20 | 7 | F | Idiopathic ASD |
| 21 | 8 | M | Tourette's ASD |
| NEUROTYPICAL CONTROLS | | | |
| 22 | 5 | F | Neurotypical |
| 23 | 6 | F | Neurotypical |
| 24 | 7 | F | Neurotypical |
| 25 | 8 | M | Neurotypical |
| 26 | 13 | M | Neurotypical |
| 27 | 7 | M | Neurotypical |
| 28 | 16 | F | Neurotypical |
| 29 | 19 | M | Neurotypical - Athlete |
| 30 | 18 | F | Neurotypical |
| 31 | 16 | M | Neurotypical |
| 32 | 17 | M | Neurotypical |

**Supplementary Table 5. Demographics of Patients with Parkinson's Disease and age-matched controls**

| Participant[1] | Gender | Age | UPDRS | Hoehn and Yahr scale | Medication (Motor/Non-Motor)[2] |
|---|---|---|---|---|---|
| PPD 1 | M | 56 | 6 | 2.5 | Yes/Yes |
| PPD 2 | F | 64 | 44 | 4 | Yes/Yes |
| PPD 3 | M | 65 | 24.5 | 2.5 | Yes/Yes |
| PPD 4 | M | 70 | 19 | 3 | Yes/Yes |
| PPD 5 | M | 59 | 29 | 3 | Yes/Yes |
| PPD 6 | F | 69 | - | 3 | Yes/No |
| PPD 7 | M | 64 | 16 | 2 | Yes/Yes |
| PPD 8 | M | 76 | 21 | 3 | Yes/No |
| PPD 9 | F | 77 | 27 | 3 | Yes/Yes |
| PPD 10 | M | 50 | - | 2 | Yes/Yes |
| PPD 11 | M | 63 | 20 | - | - |
| Elderly NT 1 | F | 65 | - | - | - |
| Elderly NT 2 | M | 68 | - | - | - |
| Elderly NT 3 | F | 57 | - | - | - |
| Elderly NT 4 | F | 76 | - | - | - |
| Elderly NT 5 | M | 77 | - | - | - |

[1]Participant Abbreviation

PPD: Patient with Parkinson's Disease

Elderly Control: age matched neurotypical control

[2] Intake of medication for motor issues / medication for non-motor issues that are known to affect motor performance

**Supplementary Table 6: Abbreviations of Body Parts**

| Body Part | Abbreviation of body part for FMR1/PPD dataset | Abbreviation of body part for Shank3 dataset |
|---|---|---|
| Pelvis | P | |
| Lumbar Level 5 | L5 | L |
| Lumbar Level 3 | L3 | L |
| Thoracic Level 12 | T12 | T |
| Thoracic Level 8 | T8 | T |
| Neck | N | |
| Head | H | H |
| Right Shoulder | RSH | RSH |
| Right Upper Arm | RUA | RUPP |
| Right Forearm | RF | RFORE |
| Right Hand | RH | |
| Left Shoulder | LSH | LSH |
| Left Upper Arm | LUA | LUPP |
| Left Forearm | LF | LFORE |
| Left Hand | LH | |
| Right Upper Leg | RUL | RTH |
| Right Lower Leg | RLL | RSK |
| Right Foot | RFoot | RFT |
| Right Toe | RT | |
| Left Upper Leg | LUL | LTH |
| Left Lower Leg | LLL | LSK |
| Left Foot | LFoot | LFT |
| Left Toe | LT | |

### *A form of Noise Cancellation with Possible Therapeutic Value*

The value of examining feedback loops, timing, and stochastic signatures across the population, can be better appreciated within the context of noise cancellation. Noise cancellation helps engineer solutions and build accommodations for PPD and other disorders of the nervous systems with tremor-related signals. Along those lines, here we investigated how removing different frequency bands from the angular speeds of the PPD affected the network dynamics. We tried different combinations of low and high limits for the frequency stop band to filter out tremor components (as reported by the literature 52 and performed *t-tests* between the resulting group and the elderly or young control groups for each body node. Then, we measured the number of nodes in which the two groups differed significantly in the NSR. We then interpolated a surface on the parameter identification data.

We report that the optimal tremor removal when comparing to the healthy young controls group is for frequencies above 19 Hz, while the optimal tremor removal when comparing to the old controls groups is for frequency band with low limits near 10 Hz. Figure 9 shows these differences.

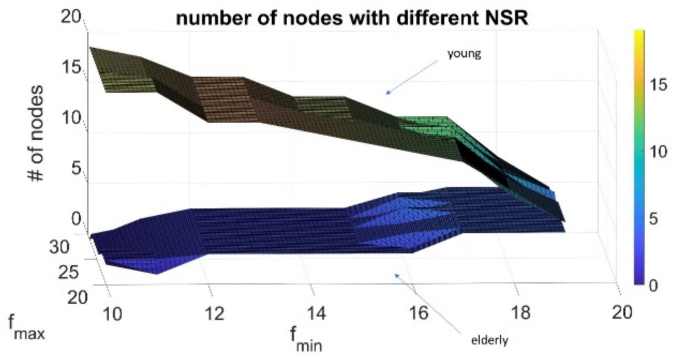


Supplementary Material Figure 1: Noise cancellation by systematic frequency removal in young *vs.* elderly participants. The two surface plots show how after removing a component of tremor from the PPD by using band stop filtering of the angular velocity for different choices of frequency, the resulting body node networks is comparable to the NSR dimension of the young and elderly controls groups. The color bar reflects the number of nodes with different NSR that results from the noise cancellation.

### *MMS Amplitude and Timings are Not Independent*

Lastly, to further understand the spatio-temporal characteristics of the MMS signal derived from the angular speed, we calculated the empirical joint distributions of the MMS inter-peak-interval times and the distributions of the MMS amplitude (peak values). We asked if they were independent. The results are shown in Figure 10. The complex patterns that emerge indicate that assuming independence between the rate of MMS activity and the MMS values is only a first approximation to a more complex mechanism that needs to be further investigated.

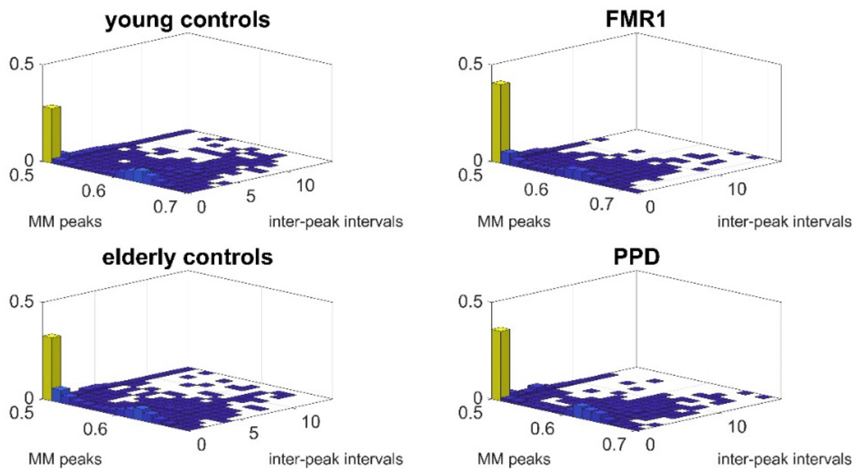


Supplementary Material Figure 2: Joint empirical distribution of MMS peak values and inter-peak-interval times.
